# GET3B interacts with the thylakoidal ALB3 and ALB4 insertases and is involved in the initial stages of chloroplast biogenesis

**DOI:** 10.1101/2024.12.05.626995

**Authors:** Uwe Bodensohn, Beatrix Dünschede, Chiara Kuhlmann, Khushbu Kumari, Roman Ladig, Christopher Grefen, Enrico Schleiff, Donna Fernandez, Danja Schünemann

## Abstract

Protein targeting and insertion into membranes are essential for cellular organization and organelle function. The Guided Entry of Tail-anchored (GET) pathway facilitates the post-translational targeting and insertion of tail-anchored (TA) membrane proteins. *Arabidopsis thaliana* has four GET3 homologues, including *At*GET3B and *At*GET3D localized to chloroplasts. These photosynthetic organelles possess complex membrane systems, and the mechanisms underlying their protein targeting and membrane biogenesis are not fully understood. This study conducted a comprehensive proteomic analysis of *get3b* mutant plastids, which displayed significant alterations. Fluorometric based complex assembly as well as CO_2_ assimilation analyses confirmed that disruption of GET3B function displayed a significant impact on photosystem II assembly as well as carbon fixation, respectively, indicating a functional role in chloroplast biogenesis. Additionally, genetic interactions were found between GET3B and the two component STIC system, which cooperates with the cpSRP pathway and is involved in the co-translational sorting of thylakoid proteins. Further, physical interactions were observed between GET3B and the C-terminus of ALB3 and ALB4 *in vitro* and the full length proteins *in vivo*, indicating a role of GET3B in protein targeting and membrane integration within chloroplasts. These findings enhance our understanding of GET3B’s involvement in stromal protein targeting and thylakoidal biogenesis.

## Introduction

Intrinsic membrane proteins (IMPs) represent an extremely diversified group that represents roughly 30 % of the cellular proteome (Wallin and Von Heijne, 1998). Tail anchored (TA) proteins share a simple type II topology with a cytosolically exposed N-terminus and a single C-terminal transmembrane domain (TMD) within the last 50 amino acids (AAs) of the polypeptide stretch. Even though they constitute 3-5% of eukaryotic IMPs (Kalbfleisch et al., 2007), TA proteins fulfill a multitude of essential biological functions like apoptosis (Jiang et al., 2014), vesicle transport (van Berkel et al., 2020) and organelle biogenesis (Sommer et al., 2013). The guided entry of TA proteins (GET) pathway facilitates the post-translational targeting of this subset of IMPs. After translation is terminated, the pretargeting complex captures the TA protein and hands it over to the central ATPase of the GET pathway, Get3 (Wang et al., 2010). Thereafter the TA protein is passed on to its cognate receptor, which functions as an insertase (Wang et al., 2014). For reasons of simplicity, we will generally refer to “Get” factors and plant specific elements will be denoted in all capital letters, in agreement with *Arabidopsis thaliana* nomenclature.

A characteristic trait of Get3 is the formation of a symmetric homodimer equipped with a static ATPase domain and a flexible α-helical domain that is sensitive to nucleotide binding. This nucleotide binding state induces the formation of a hydrophobic groove or chamber capable of binding TA substrate (Mateja et al., 2009; Bozkurt et al., 2009, Mateja et., 2015, Barlow et al., 2023). In a cellular context, Get3 also has a secondary function in eukaryotes by acting as an ATP-independent holdase under energy limiting conditions and oxidative stress (Powis et al., 2013; Voth et al., 2014). Get3 and its orthologues TRC40/ArsA are highly conserved in all domains of life and have diversified into three clades, namely clade a, clade bc and clade d (Xing et al., 2017; Barlow et al., 2023). In contrast to metazoans and fungi, which solely contain clade a Get3 homologues, photosynthetic organisms contain multiple Get3 homologues from all clades (Xing et al., 2017; Bodensohn et al., 2019; Barlow et al., 2023). The three clades differ in their amion acid (AA) composition, which results in different domain architectures. In the majority of the cases clade a proteins have a cytosolic and clade bc an organellar localization (Xing et al., 2017; Bodensohn et al., 2019). Depending on the prokaryotic or eukaryotic origin of a photosynthetic organism, clade d proteins have a cytosolic or organellar localization, respectively (Barlow et al., 2023).

*Arabidopsis thaliana* has four Get3 homologues, namely *At*GET3A, *At*GET3B, *At*GET3C and *At*GET3D localized to cytosol, chloroplasts, mitochondria and chloroplasts, respectively (Duncan et al., 2013; Xing et al., 2017; Bodensohn et al., 2019; Barlow et al., 2023). Loss of function mutations in any of the clade a or clade bc orthologues renders the resulting plant largely indistinguishable from the wild-type (Xing et al., 2017). This is also the case in yeast and nematodes in which the deletion of Get3 leads to conditional growth defects (Schuldiner et al., 2008; Tseng et al., 2007). In mammals however, the disruption of TRC40 is embryo lethal (Mukhopadhyay et al., 2006).

In photosynthetic eukaryotes, plastids constitute one of the most complex systems in terms of protein sorting. 95 % of their proteome is encoded by nuclear genes and precursor proteins are synthesized on cytosolic ribosomes to be imported post-translationally. This is facilitated by two multimeric translocons embedded in the outer (TOC) and inner (TIC) envelope that form a continuum from the cytosol, through the intermembrane space (IMS) into the stroma (Liu et al., 2023). Once arrived in the stroma, precursor proteins can be directed to numerous translocation machineries imbedded within the inner envelope or thylakoid membrane. This leads to six different sub organellar localizations that each can individually be accessed with different sets of protein targeting networks (Jarvis and López-Juez, 2013). Taking the pivotal role of plastids into account, the targeting and translocation networks must operate with high fidelity to ensure proper organellar biogenesis, repair and functionality. This is reflected by the deficiencies in plant fitness that arise when components of these are disrupted (Bauer et al., 2000; Amin et al., 1999; Pilgrim et al., 1998; Sundberg et al., 1997).

One intensively studied stromal targeting pathway is the post-translational cpSRP pathway, which targets a single population of IMPs, namely light harvesting chlorophyll binding proteins, to the thylakoid membrane (Schuenemann et al., 1998; Ziehe et al., 2018). CpSRP is composed of cpSRP54 (henceforth referred to as SRP54), the plastidic homologue of a highly conserved subunit of the cytosolic SRP, and a plant specific targeting factor, cpSRP43 (SRP43 from here on). SRP54 can also interact with the plastidic 70S ribosome and mediates co-translational transport of substrate proteins to its respective insertase, ALB3 and the SecYE translocon (Hristou et al., 2019; Moore et al., 2003).

The terminal insertase ALB3 belongs to the Oxa1 superfamily of membrane protein biogenesis factors which also includes the ER resident Get1 (Anghel et al., 2017). In plants, an additional homologue of ALB3 exists, ALB4 (Gerdes et al., 2006). ALB4 is involved in the assembly and stabilization of ATP synthase intermediates (Benz et al., 2009). Furthermore, subsequent analyses displayed that ALB4 works at least partially together with ALB3 and participates in the biogenesis of a subset of thylakoid membrane proteins. (Trösch et al., 2015).

ALB4 also interacts with STIC2, a protein identified in a genetic screen for the suppressors of chlorotic *tic40*, an *A. thaliana* mutant lacking the chloroplast inner envelope protein TIC40. The STIC system is described as a two component network composed of STIC1 (ALB4) and STIC2, which acts together in a shared pathway with cpSRP54 and cpFtsY (Bédard et al., 2017). STIC2 shares homology with the bacterial YbaB,(Link et al., 2008). Recently, STIC2, was identified as a factor associated with plastidic ribosomes translating the photosystem (PS) II reaction center protein D1, is likely involved in co-translational sorting of other plastid-encoded multi-pass IMPs (Stolle et al., 2024).

In a previous study, we provided evidence that the *At*GET3B can selectively bind SECE1 and shows a genetic interaction with components of the cpSRP system. Furthermore, mutant plants showed altered levels of photosynthesis-related proteins (Anderson et al., 2021). This prompted us to investigate if GET3B interacts with constituents of the cpSRP and STIC systems, with an emphasis on their cognate receptors ALB3 and ALB4, respectively. We hypothesized that GET3B might play a crucial role in the targeting and membrane integration of specific proteins within chloroplasts, thereby contributing to the biogenesis and functionality of chloroplast membranes. To address this hypothesis, we employed a combination of biochemical, molecular, fluorometric and genetic approaches to provide evidence that GET3B physically interacts with the C-terminus of ALB3 as well as ALB4 and genetically interacts with components of the STIC and cpSRP pathways, and that its disruption has an effect on chloroplast biogenesis and function.

## Materials and Methods

### Plant materials and plant growth

*Arabidopsis thaliana* seeds were surface sterilized and sown on 0.8% (w/v) agar plates containing ½ strength Murashige and Skoog media (Caisson Labs), 50mM MES-KOH, pH 5.7, and 1% (w/v) sucrose. Plants used for fluorometric measurements were germinated and cultivated on MS plates without sucrose. Following 2-3 days of stratification at 4°C, the plates were exposed to 100 μE m^−2^ s^−1^ of light provided by cool white (4100K) fluorescent bulbs on a 12 h light/12 h dark cycle for 7-14 days at 22°C, at which point seedlings were transplanted to a soil-less mix (Fafard Germinating Mix, Sungro Horticulture), watered with 10% Hoagland’s solution (Hoagland and Arnon, 1950), and grown under the same conditions as the plates.

Columbia-0 ecotype was used as wild-type for all pictures and measurements in this study. Seeds for the T-DNA insertional mutants were acquired from the Arabidopsis Biological Resource Center (https://abrc.osu.edu/). Mutant alleles previously characterized as resulting in loss of function were used for the genetic crosses: *get3b-2* (SALK_017702C, Xing et al., 2017), *cpsrp54-3* (WiscDsLox289_292B14, Yu et al., 2012), *alb4-1* (SALK_136199, Bédard et al., 2017), *stic2-3* (SALK_001500, Bédard et al. 2017) and *stic2-4* (WiscDsLox445D01, Bédard et al. 2017). The two *stic2* alleles behaved similarly; therefore, we refer to *stic2* mutants without the allele number in other sections of the paper. Lines carrying the mutant alleles were identified by PCR genotyping using the genotyping primers listed in Supplemental Table S2.

### Generation of transgenic plants

Generation of the constructs used for moderate overexpression and genetic complementation experiments was previously described (Anderson et al. 2021). The *ProUBI10:GET3B-STREP* construct contains the promoter for the *UBIQUITIN 10* gene (Norris et al., 1993), genomic sequence for the coding region of *GET3B*, sequence encoding a C-terminal 1X STREP tag, and the terminator of the *NOS* (*nopaline synthase*) gene. For the *ProUBI10:GET3B(D124N)-STREP* construct, the D124N mutation was introduced into the *ProUBI10:GET3B-STREP* construct by site-directed mutagenesis. The Arabidopsis plants transformed for the genetic complementation experiments were homozygous for the *alb4* and *get3b* alleles, and heterozygous for the *srp54* allele. Plants transformed for moderate overexpression studies were homozygous for either the *alb4* or *stic2-4* allele. The plant transformation procedure was previously described (Anderson et al. 2021), and transformants were selected in the T1 generation based on gentamycin resistance. Transformants containing mutant alleles as well as the introduced transgene were identified by PCR genotyping using primers listed in Supplemental Table S2. Genetic complementation was assessed in the T2 generation based on the phenotype of plants that were homozygous for *alb4*, *get3b* and *srp54* and carried the introduced transgene. The effects of moderate overexpression in *alb4* or *stic2* homozygous backgrounds were assessed in either the T2 or T3 generations.

### Rosette measurements

For measurements of rosette diameters and chlorophyll content, plants with specific genotypes were grown for approximately 4 weeks under the conditions described above. Rosette diameter was determined by measuring across the rosette at the point where the distance was largest. For chlorophyll measurements, 120 seedlings were collected for each genotype or transgenic line. These were pooled and weighed to estimate the freshweight. Chlorophyll was extracted with 80% acetone in 2.5 mM HEPES-KOH (pH 7.5) as described previously (Hackett et al., 2017) and chlorophyll amounts were calculated from absorbances measured on a spectrophotometer, as described in Wellburn (1994). Pairwise Student’s *t* tests (two-tailed) were used to compare means of rosette diameter and ANOVA to compare the chlorophyll concentration.

### Chloroplast isolation

Chloroplasts from leaf tissue of *Arabidopsis thaliana* were isolated according to (Bodensohn et al., 2020).

### Pulse amplitude modulation (PAM) measurements

Seedlings cultivated as described above were first dark adapted for 15 min before being placed in the FluorCam 800-C (Photon systems instruments) for measurements. Initially F_0_ was recorded for 5 sec with a measuring beam. Thereafter F_M_ was initiated with a saturating flash at 4000 µE. After a relaxation phase of 17 sec actinic light at 150 µE was utilized to assess the photosynthetic adaptation (non-photochemical quenching NPQ) for 5 pulses every 12 sec. This was followed by another measurement in the dark for 3 pulses.

### Infrared gas analyzer measurements (IRGA)

The gas exchange rate of 4week-old plants was measured using a LI-6400XT portable photosynthesis system (Li-Cor Biosciences Inc.). Five individual plants, grown under short-day conditions (8-h light provided by cool white fluorescent bulbs, 100 µM m^−2^s^−1^, 22 °C; 16-h dark, 20 °C) were used for each experiment. For the light-dependent tests, the whole-plant chamber was set up with the following parameters: 200 µmol/sec flow rate, 20◦C block temperature, 50–70% relative humidity, and 400 µmol/mol [CO2]. For differential light conditions, the plants were dark-exposed (0 PAR) for 30 min and then light-exposed (400 PAR) for 30 min followed by another dark (0 PAR) phase for 30 min. The data was logged every 30 sec. The LI-6400XT portable photosynthesis system (Li-Cor Biosciences Inc.) generates the raw data for the rate of transpiration over time with a default value for the area of the plants. Following the gas exchange rate measurement, ImageJ was used to determine the plant’s area (Schindelin et al., 2012). The rate of transpiration measurement was automatically standardized on the plant area by adding this value to the raw data from the LI-6400XT device. Furthermore, the time scale was modified in accordance with the experiment. For instance, the graph’s x-axis can be modified to show 30 minutes on the negative scale and 60 minutes on the positive scale once the time point 0 is set at 400 PAR light. Data presented are means of at least three leaves from individual plants for each experiment. Experiments were repeated at least three times.

### Plasmids and plasmid construction for protein expression

For the overexpression of GST-tagged mature full-length GET3b (GST-Get3b), the corresponding sequence encoding amino acids (aa) 68-411 was cloned into pGex-4T-3 using the BamHI/SalI restriction sites. The length of the chloroplast transit sequence was predicted by the TargetP-2.0 server (https://services.healthtech.dtu.dk/services/TargetP-2.0/).

Constructs for the expression of His-tagged C-terminal regions of ALB3 corresponding to aa 350-462 (Alb3C-His) and ALB4 corresponding to aa 334-499 (Alb4C-His) were described previously (Ackermann et al., 2021).

### Expression and purification of recombinant proteins

His-tagged proteins were expressed in *E. coli* strain Rosetta2(DE3). Cells were cultivated in LB medium at 37 °C to an optical density between 0.6 and 0.8. After induction with 1 mM of isopropyl-β-D-thiogalactopyranosid (IPTG) the cells were grown for additional 3 hours at 30 °C. Collected cells were resuspended in washing buffer (20 mM HEPES, 300 mM NaCl, 40 mM imidazole, 2 mM DTT, pH 8.0). Cells were disrupted by sonification and after centrifugation, the supernatant was either loaded onto a nickel-nitrilotriacetic acid column (GE Healthcare) or incubated with nickel-nitrilotriacetic acid resin (Qiagen). After washing with washing buffer the fusion constructs were eluted with elution buffer (20 mM HEPES, 300 mM NaCl, 250 mM imidazole, 2 mM DTT, pH 8.0). Subsequently, the imidazole was removed from the buffer using PD-10 columns (GE Healthcare). The protein was kept on ice and used directly or was stored at −20°C until usage.

GST-Get3b was expressed as described above with the following modifications. Collected bacterial cells were resuspended in 1x PBS buffer (140 mM NaCl, 2.7 mM KCl, 10 mM Na_2_HPO_4_, 1.8 mM KH_2_PO_4_, pH 7.3) before sonication. Proteins were purified using glutathione-sepharose (GE Healthcare) and eluted with 10 mM reduced glutathione in 1xPBS buffer. The buffer was exchanged to 1xPBS using PD-10 columns (GE Healthcare).

### Pull-down assay

*In vitro* pull-down analyses were performed using 50 μg of GST-Get3b and either 20 μg of Alb3C-His or 20 μg of Alb4C-His in a total volume of 120 μl of 1x PBS. Negative control reactions were conducted using GST instead of GST-Get3b or the His-tagged constructs alone. The samples were incubated for 10 min at room temperature. Subsequently, 100 μl of a 50% slurry of glutathione resin (equilibrated in 1x PBS) were added and incubated for 30 min at room temperature. The mixture was transferred to Wizard® Mini Columns (Promega) and centrifuged briefly. The column was washed with 5 ml of 1xPBS buffer using a syringe. After a brief centrifugation of the column the proteins were eluted with 30 μl of 10 mM reduced glutathione in 1xPBS buffer. The load as well as the eluted samples were analyzed by SDS-PAGE and proteins detected by Coomassie staining or Western blotting.

### Construction of split-ubiquitin plasmids and the split-ubiquitin yeast two-hybrid assays

For the split-ubiquitin yeast two-hybrid assay the coding sequence for mature full-length GET3b (aa 68-411) was cloned into pADSL-Nx using the BamHI/SalI restriction sites. All other constructs have been described previously (Bals et al., 2010; Pasch et al., 2005). The split-ubiquitin assay was done according to (Pasch *et al*, 2005).

### MS/MS analysis

For in-solution mass spectrometry, proteins were digested (León et al., 2013; ISD:SDC; AP&PT) and peptides were purified by C18 STAGE-TIPS (Rappsilber et al., 2007). Peptides were analyzed using an ultra-HPLC Proxeon EASY-nLC 1000 system coupled online to Q Exactive Plus mass spectrometer (Thermo Fisher Scientific). Reversed-phase separation was performed using a 30 cm analytical column (100 μm diameter; DNU-MS (Novak) packed in-house with Reprosil-Pur 120 C18-AQ, 2.4 µm). Mobile-phase solvent A consisted of 0.1% formic acid and 4% acetonitrile in water, and mobile-phase solvent B consisted of 0.1% formic acid in 80 % acetonitrile. The flow rate of the gradient was set to 200 nl/min. A 70-min gradient was used (0–40% solvent B within 40 min, 40–100% solvent B within 10 min, 100% solvent B for 10 min, 100–0% solvent B within 5 min and 0% solvent B for 5 min).

Data acquisition was performed with the ddMS2 method with the following configuration: For the MS scans, the scan range was set to 250–2,000 m/z at a resolution of 70,000, and the automatic gain control (AGC) target was set to 1 × 10^6^. For the MS/MS scans, Top 13 ions were chosen, the resolution was set to 35,000, the AGC target was set to 1 × 10^5^, the precursor isolation width was 2 Da and the maximum injection time was set to 80 ms. The HCD normalized collision energy was 27%. MaxQuant was used to analyze the LC-MS/MS data (ver 1.5.5.; Cox and Mann, 2008) which allowed qualitative and quantitative analysis. The UniProt reference *Arabidopsis* database (UP000006548) was used for the identification of proteins. Default settings for fixed modifications were used, Dynamic modifications were set: Oxidation for M and Deamidation for NQ. Contaminants were included for peptide detection of a minimum length of 6 amino acids. FDR threshold was set to 0.01. Perseus was used to transform the quantitative data and perform a statistical analysis (Tyanova and Cox, 2018). The default settings for a Student’s t-test were utilized for the analysis. For this, independent triplicates corresponding to roughly 80 µg of plastid protein isolated from 8 pots of *Arabidopsis thaliana* lawn from wild-type and *get3b* plants were compared.

### Accession numbers

The following Arabidopsis genes were investigated in this study: AT3G10350 (GET3B), AT5G03940 (cpSRP54), AT1G24490 (ALB4/STIC1), and AT2G24020 (STIC2).

## Results

### Proteomic characterization of the *get3b* mutant

For the GET3B locus, three T-DNA insertion lines are publicly available (Xing et al., 2017). We decided to analyze the *get3b-2* mutant (from here on *get3b*) with an insertion in the fourth intron of the allele. The line was homozygous for the mutation and had no transcript nor protein detectable by RT-PCR or western blot, respectively (Figure 1A-D).

**Figure 1.**
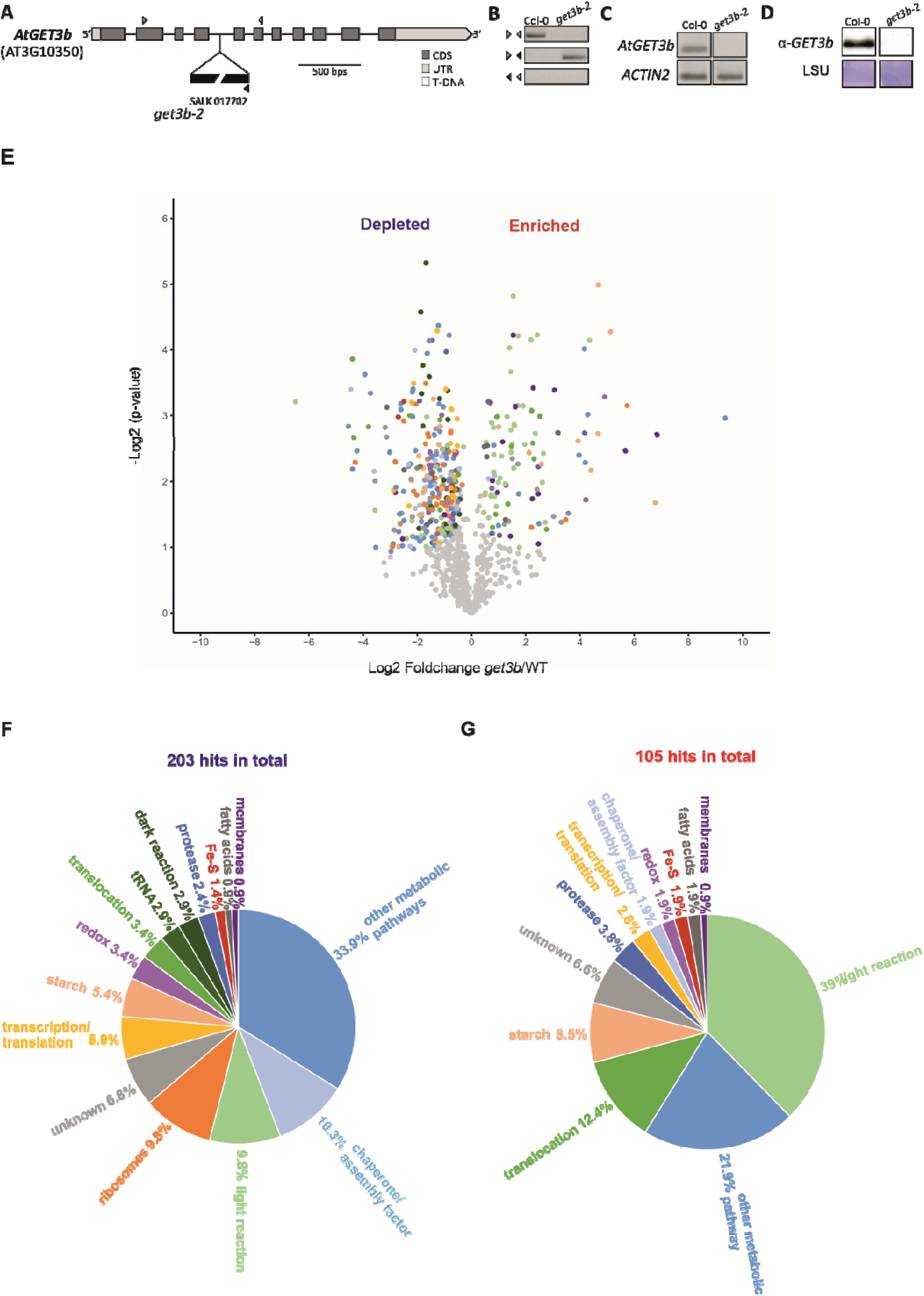
The *get3b* mutation and the downstream effects on the chloroplast proteome. **(A)** Gene model of *GET3B* and the corresponding T-DNA insertion line analyzed in this study. **(B)** Genotyping via PCR of *get3b*. The locations of the used primers are indicated in A as arrowheads. **(C)** Semi RT-PCR to validate the missing transcript. **(D)** Western blot analysis of total protein lysates. Wild-type and *get3b* plastids were isolated and subjected to LC-MS/MS followed by label free quantification. **(E)** Statistical analysis of proteinaceous hits that were significantly effected in *get3b* chloroplasts (see Supplemental table S1 for more details). The significantly depleted **(F)** and significantly enriched **(G)** proteinaceous hits were categorized according to their cellular function. The color coding in E corresponds to those in F and G.

To analyze the overall effect of the *get3b* mutation on chloroplast function, we examined *get3b* chloroplasts on a proteomic level. By using wild-type and *get3b* plastids, a label-free technique could be employed in a quantitative manner as previously described (Vermeulen et al., 2008). For this, we isolated plastids of the respective lines from 21-day-old *A. thaliana* seedlings and subjected these to LC-MS/MS.

The examination revealed a total of 999 proteinaceous hits. After subtraction of contaminants, 203 were classified as significantly depleted and 105 as enriched in *get3b* (Figure 1E). Intriguingly, not only nuclear encoded proteins, but also plastome encoded proteins were significantly affected. These covered a wide range of biological processes and were manually curated and categorized according to their cellular functions (Figure 1F, G). The majority of the significantly depleted hits (33.9 %) belonged to the category *other metabolic pathways*, which included components of the AA synthesis machinery as well as the post-translational modification machinery. Apart from that, *chaperone/assembly factors* (10.3 %), the translational machinery (9.8 % *ribosomes* plus 5.9 % *transcription/translation*), the photosynthetic machinery (9.8 % *light reaction* plus 2.9 % *dark reaction*) were mostly affected. Constituents of *starch synthesis* (5.4 %), the plastid *redox* regulation and *translocation* (3.4 % each), tRNA (2.9 %), *proteases* (2.4 %), *Fe-S cluster* synthesis (1.4 %), *fatty acids* and *membranes* (0.9 % each) were also depleted to a significant extent in *get3b* (Figure 1F). Interestingly, one entire paralogous set of nuclear encoded proteins of the oxygen evolving complex (OEC; Allahverdiyeva et al., 2013) was reduced to a significant level. Furthermore, several components of the electron transport chain were also decreased. Additionally, soluble proteins involved in the Benson-Calvin cycle like several subunits of ribulose-1,5-bisphosphat-carboxylase/-oxygenase (RuBisCo) and their assembly factors were also negatively affected as well as STIC2, a factor involved in stromal targeting networks (Bédard et al., 2017). Strikingly, GET3B was detectable even though it was significantly depleted (Supplemental table S1), suggesting that the T-DNA insertion might lead to a knock-down rather than a knock-out of the respective gene.

The majority of the significantly enriched hits were attributed to the *light reaction* of photosynthesis (39 %), followed *by other metabolic pathways* (21.9 %) and components of *translocation* machineries (12.4 %). Additionally, *starch* synthesis (8.5 %), *proteases* (3.8 %), constituents involved in *transcription/translation* (2.8 %), *chaperone/assembly factors, redox* regulation, *Fe-S* cluster biogenesis and *fatty acid* synthesis (1.9 % each) as well as elements involved in membrane modulation (0.9 %) were elevated to a significant extent (Figure 1G). Strikingly, nearly all LHCPs of PSII and a few of PSI were enriched in *get3b*, while the respective cpSRP targeting machinery was unaffected (Supplemental table S1).

The comparative proteomic data displayed that the loss or reduction of GET3B has a pervasive effect on the proteome of plastids. Interestingly, the photosynthetic apparatus seemed to be affected the most.

### *get3b* mutation affects de novo assembly of photosystem II

The data from the comparative proteomics indicated that the light reaction as well as assembly factors of the photosynthetic machinery, like YCF3 and CRR6, were affected in *get3b* (Figure 1E-G, supplemental table S1). This prompted us to examine a possible involvement of GET3B function in photosystem assembly. In order to address this, we employed PAM measurements during de-etiolation of *A. thaliana* seedlings. These measurements exploit the fluorescence solely emitted by PS II, since PS I is a fluorescence quencher (Brettel, 1997). We utilized Fv/Fm values as a read out for PS II functionality. By employing this during de-etiolation we were able to assess the extent of de novo formation of PS II. For this, seedlings were germinated and cultivated *in vitro* on Murashige Skoog (MS) medium (Murashige and Skoog, 1962) for 4 days in the dark and in the morning of the fifth day illuminated normally. The course of de-etiolation and concomitant de novo PS II assembly was encompassed by PAM measurements. This occurred 1, 7 and 24 hours after the initial illumination.

When cultivated without a carbon source, *get3b* was significantly hampered in the assembly of PSII in comparison to wild-type after 7 and 24 hours (Figure 2A; top panel). In the presence of a carbon source, this difference was only apparent after 24 hours and not to such a high degree of significance, indicating that PS II functionality was still hindered in the absence or reduction of GET3B when grown heterotrophically (Figure 2A; bottom panel).

**Figure 2.**
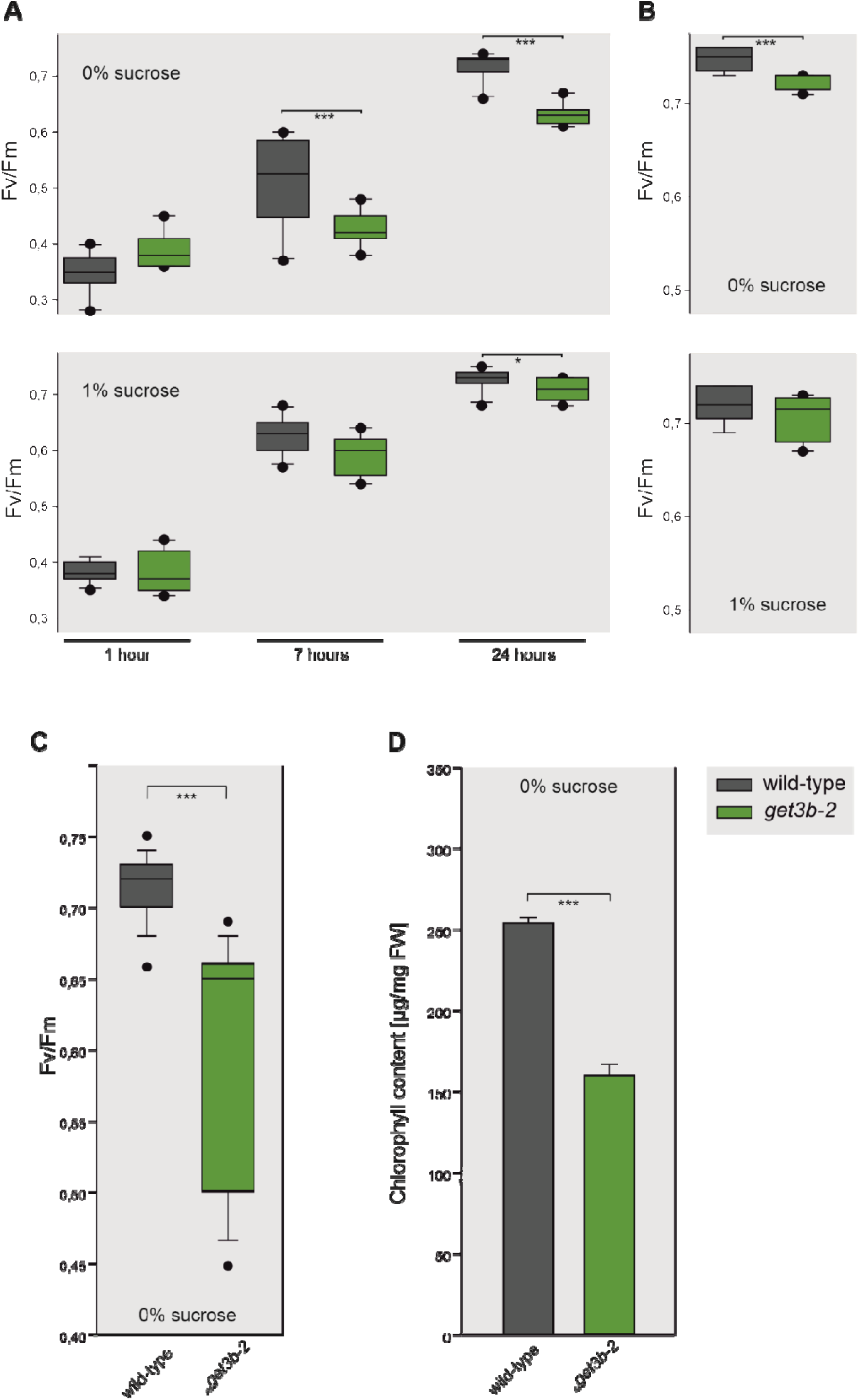
GET3B function is involved in the assembly of photosystem (PS) II. **(A)** Seedlings were germinated and cultivated under etiolating conditions for 5 days on 0% sucrose (top) or 1 % sucrose (bottom) for 5 days before being normally illuminated and subjected to pulse amplitude modulation (PAM) measurements. The Fv/Fm ratio (maximum quantum yield) was determined after 1, 7, 24 hours to monitor PS II assembly. **(B)** Seedlings were grown under normal illumination on 0 % sucrose (top) and 1 % sucrose (bottom). After 5 days these were subjected to PAM measurements. **(C)** Seedlings were cultivated under normal conditions for 4 days and either subjected to PAM measurements or **(D)** used for chlorophyll isolation. Statistical analyses for (A) and (B) were performed using the Student’s t-test, for three independent experiments with 40 seedlings per experiment. Statistical analyses for (C) were performed with ANOVA for 15 independent experiments with 40 seedlings per experiment as well as (D) for 3 experiments with 120 seedlings per experiment.

As a control we grew the same plates for five days without etiolation and analyzed these fluorometrically. When cultivated autotrophically, *get3b* was still significantly obstructed in PS II assembly (Figure 2B; top panel). Conversely, when grown in the presence of a carbon source, no difference was detectable with PAM measurements (Figure 2B; bottom panel).

Being able to observe significant differences after five days of growth under normal irradiation, prompted us to analyze 4-day-old seedlings under regular illumination in the absence of sucrose. The corresponding PAM measurements mirrored the previous assessment and confirmed an impaired PSII assembly in *get3b* (Figure 2C). Consistently, *get3b* also exhibited a significant reduction in chlorophyll content (Figure 2D).

Taken together these data indicated that, in *get3b* mutants, not only the initial stages (up to 24 hours) of de novo assembly of PS II were affected but also the later stages (up to 5 days). Additionally, the reduced chlorophyll content of *get3b* illustrated the obstruction of the photosynthetic machinery during early plant development.

### *get3b* mutation affects CO_2_ assimilation

The proteomic analysis also suggested that components of the dark reaction were affected by the *get3b* mutation. Interestingly, not only three of four paralogous genes of the nuclear encoded small subunits but also the plastom encoded large subunit of RuBisCo were affected by the reduction or absence of GET3B. Additionally, RuBisCo assembly factors like RbcX, CPN60A1, CPN60B1 and CPN10-2 were significantly affected in *get3b* (Supplemental table S1). In total, these constitute six out of nine components necessary to synthesize RuBisCo *in vitro* (Aigner et al., 2017). In order to address this matter, we analyzed the CO_2_ assimilation of *A. thaliana* seedlings by Infra-red gas analyzer (IRGA; Toro et al., 2019). The measurements revealed that *get3b* assimilated significantly less CO_2_ (Figure 3). During the measurements we included a complementation line that was previously used (Xing et al., 2017). The complementation line was able to rescue the *get3b* assimilation phenotype and restore the levels above those of the wild-type (Figure 3).

**Figure 3.**
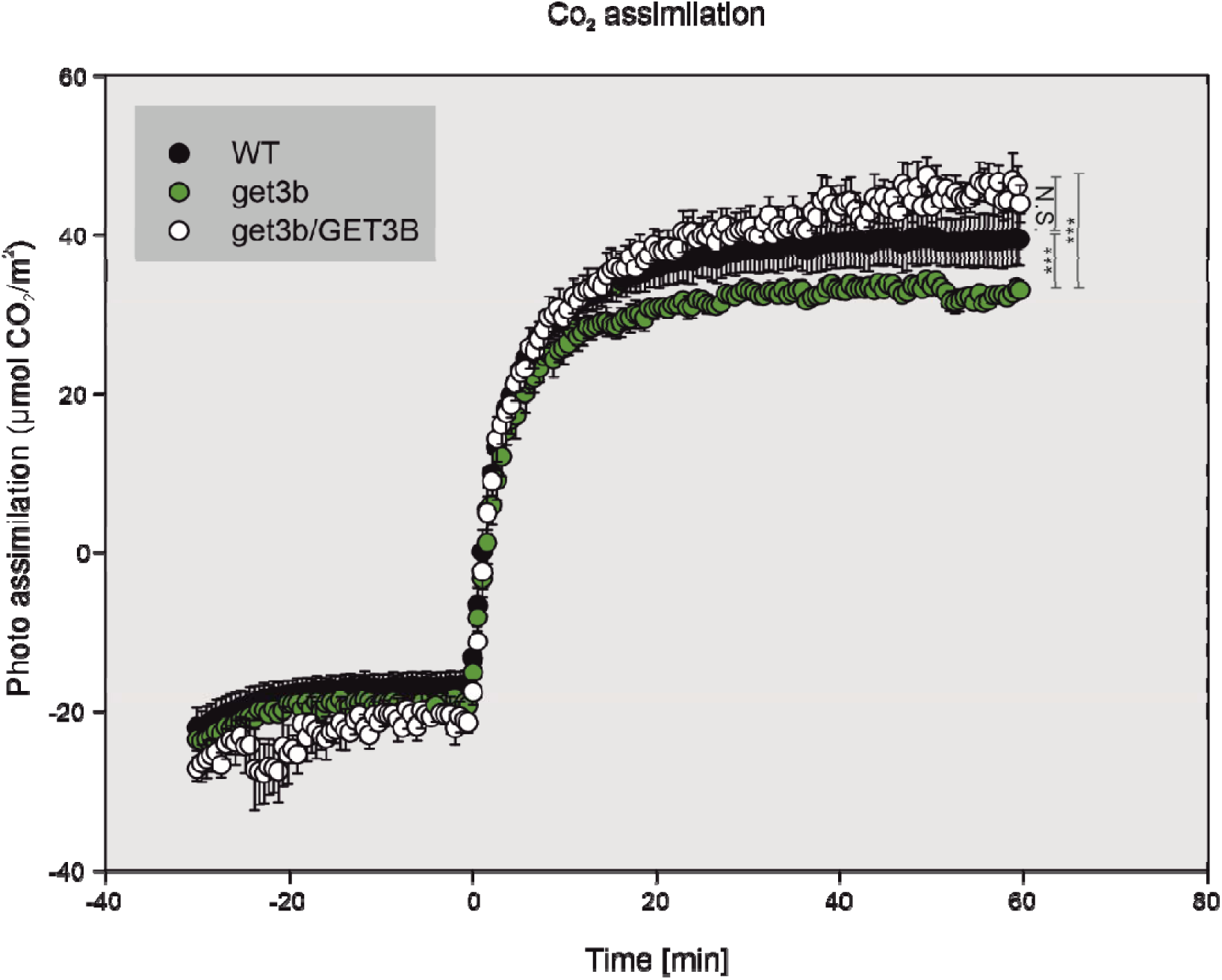
GET3B function is involved in CO_2_ assimilation. 4-week-old plants of the depicted lines were subjected to infrared gas analyzer (IRGA) measurements. Data points represent the mean of 4 independent measurements. Asterisks denote significant differences (one way anova ***P<0.001). N.S. : not significant

This and the previous data support the results of the proteomic analysis, that GET3B is required for an efficient light as well as dark reaction of photosynthesis.

### GET3B interacts genetically with components of the STIC pathway (ALB4 and STIC2)

Previously we had shown the negative -synergistic effect of combining the *get3b* and *srp54* alleles (Anderson et al., 2021). A functional interaction between the two factors also became apparent from our proteomic analysis, in which *get3b* displayed a significant enrichment of cpSRP substrates (Figure 1, supplemental table S1). This could be due to a form of compensation for the absence or reduction of GET3B function or the reduction of some stromal proteases which we could also observe in the proteomic data (Supplemental table S1). Additionally, the significant depletion of STIC2 in *get3b* prompted us to analyze the connection between the GET3B-, cpSRP- and the STIC-components.

Hence, we utilized the existing single mutants *srp54* (Amin et al., 1999), *stic2* (Bédard et al., 2017), *alb4* (Gerdes et al., 2006) and *get3b*, and double mutants *srp54 get3b* (Anderson et al., 2021), *alb4 srp54,* and *alb4 stic2* (Bédard et al., 2017), and combined mutant alleles to create the double mutants *alb4 get3b* and *stic2 get3b*, and triple mutants *alb4 stic2 get3b* and *alb4 srp54 get3b* (Figure 4). To assess the phenotypic consequences of these genetic combinations, we measured the rosette diameter as an indicator of overall plant fitness. Unlike the *get3b*, *alb4*, *stic2* single mutants and *alb4 stic2* double mutant, the *srp54* single mutant and *alb4 get3b*, *stic2 get3b* double mutants displayed a significantly smaller rosette diameter compared to the wild-type (Figure 4A-F, H, I, M). Strikingly, this phenotypical trait was further exacerbated in the *srp54 get3b* double mutant and *alb4 stic2 get3b* triple mutant (Figure 4G, J, M). These results illustrate an additive or synergistic interaction between GET3B and STIC components. Notably, the *get3b* allele in a *srp54* background showed a more severe phenotype than in an *alb4 stic2* background, suggesting a stronger functional interdependence between GET3B and SRP54.

**Figure 4.**
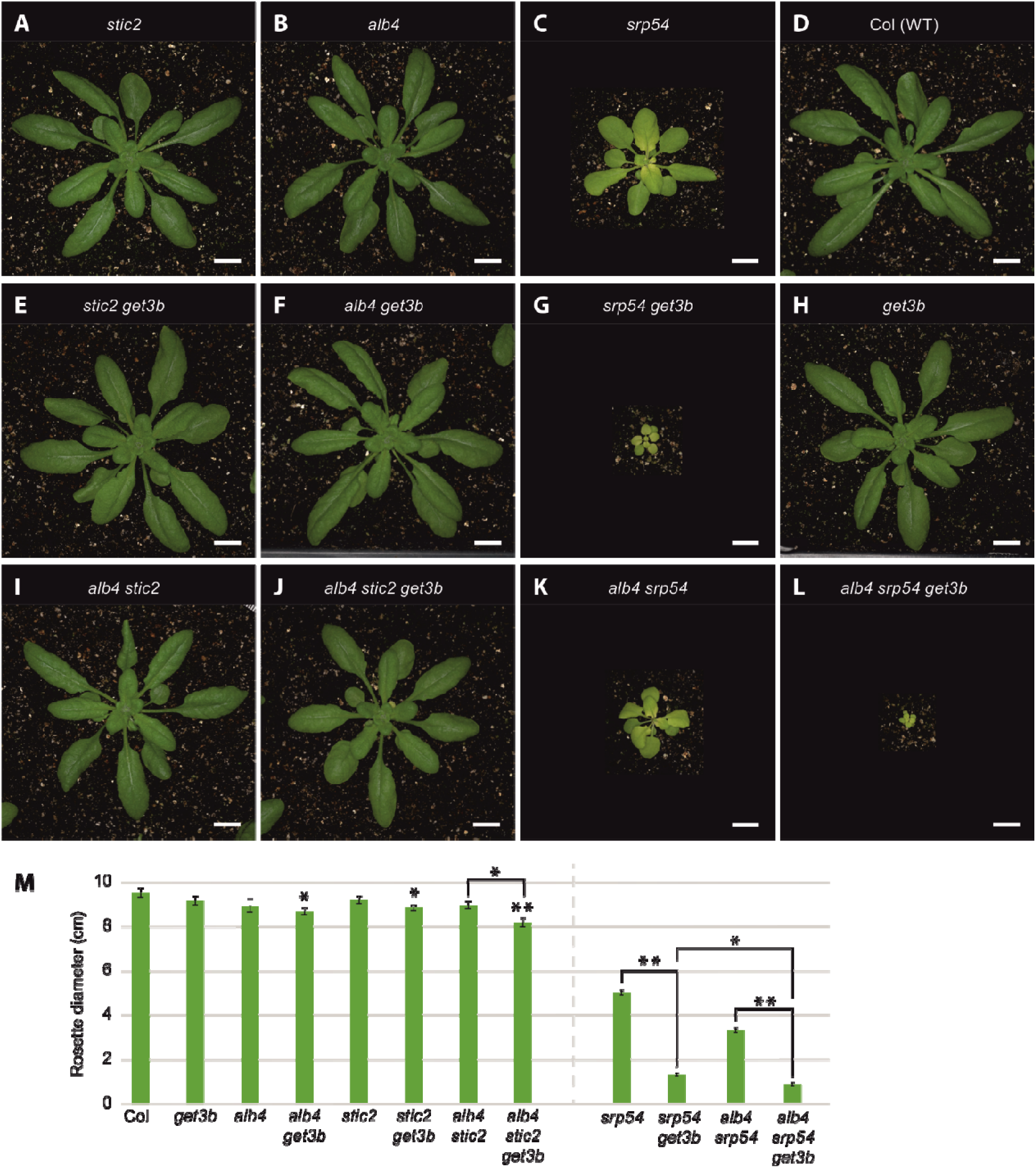
Genetic interactions are seen when mutations in *GET3B* are combined with mutations affecting *ALB4* and/or *STIC2*. Plants were grown in 12-h light/12-h dark conditions for 4 weeks. Top row: Images of **(A)** *stic2*, **(B)** *alb4* **(C)** *srp54,* **(D)** Columbia wild-type plants. Middle row: Phenotypes when *get3b* mutations were introduced: **(E)** *stic2 get3b*, **(F)** *alb4 get3b*, **(G)** *srp54 get3b*, **(H)** *get3b* plants. Bottom row: **(I)** *alb4 stic2,* **(J)** *alb4 stic2 get3b*, **(K)** *alb4 srp54*, **(L)** *alb4 srp54 get3b* plants. Black areas were added so that plants would be shown at equal magnification. Bars= 1 cm. **(M)** Rosette diameters of plants with different genotypes in the absence (left) or presence (right) of *srp54* mutations. Data are presented as means ± SE (n= 2-10 individual plants). Asterisks designate significant differences (Student’s *t* test, \**P*<0.05, \*\**P*<0.01) from wild-type or between the samples indicated with brackets.

In order to better understand this interdependence as well as the effect of the *get3b* allele on plant fitness, we analyzed the *alb4 srp54 get3b* triple mutant. While *alb4 srp54* plants were smaller than *srp54* plants (Figure 4K, M), the addition of the *get3b* allele resulted in a further reduction in the measured rosette diameter (Figure 4L, M). This demonstrated that the combination of the *alb4*, *srp54* and *get3b* alleles resulted in an additive detrimental effect on plant fitness.

### The ATPase function of GET3B is responsible for the growth phenotype

In order to investigate the restorative effects of GET3B expression in the *alb4 srp54 get3b* triple mutant, we complemented the triple mutant by transfecting the plants with a construct previously used (Anderson et al., 2021). In the T2 generation, 12 independent lines were identified carrying the *ProUB10:GET3B-STREP* transgene. Data are shown for 3 randomly chosen lines (Figure 5B-C, G). The rosette diameters of the three lines were significantly smaller (lines 2,3) as well as larger (line 1) than the *alb4 srp54* double mutant (Figure 5A-C, G). However, in all of the lines, rosette diameters were more than three times larger than the *alb4 srp54 get3b* triple mutant, signifying that the transgene was able to rescue the triple mutant phenotype (Figure 5G, H).

**Figure 5.**
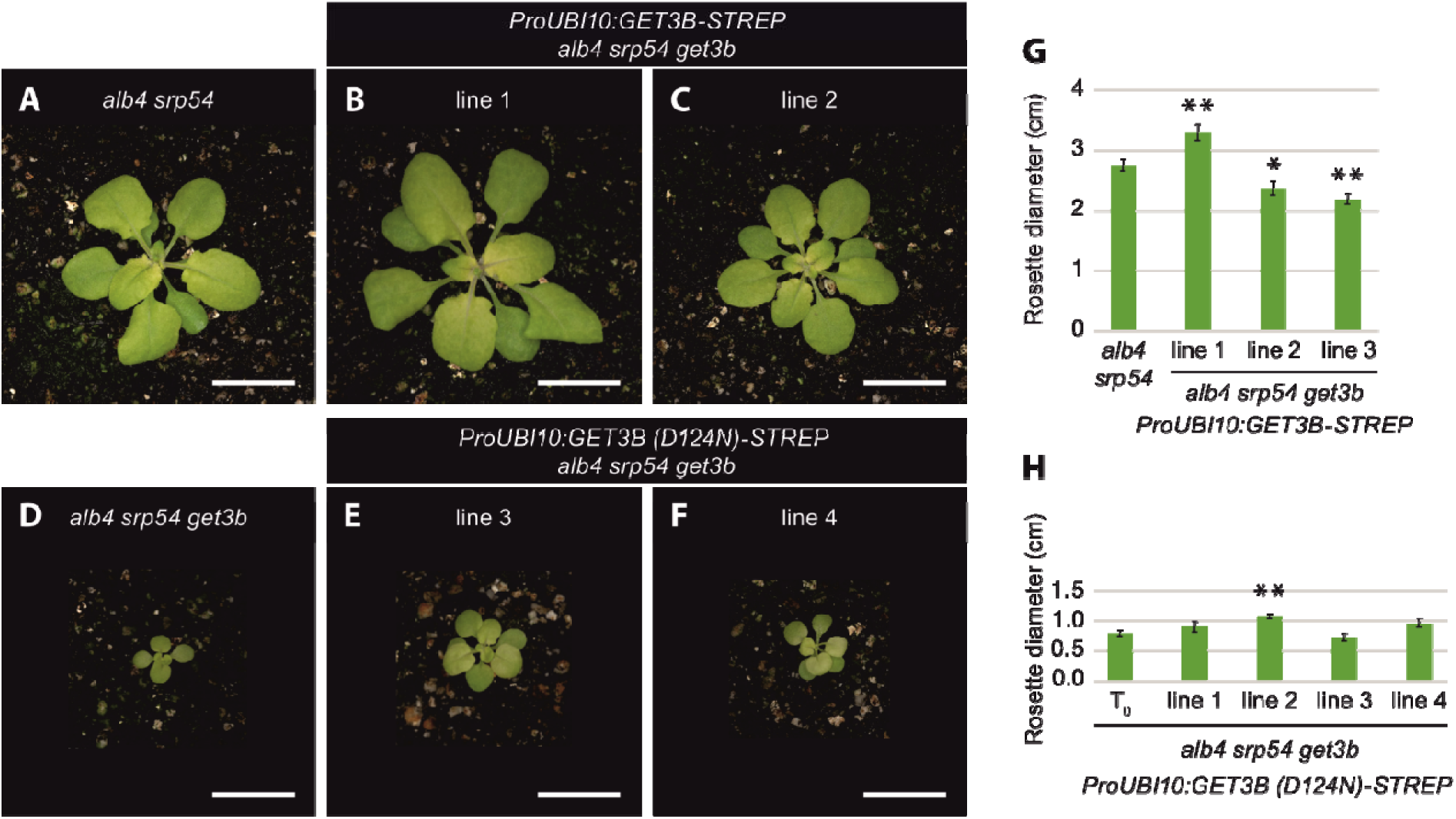
Transgenes encoding wild-type GET3B can complement alb4 srp54 get3b triple mutants, but not transgenes encoding GET3B with a D124N mutation. Plants were grown in 12-h light/12-h dark conditions for 4 weeks. Top row: Images of (A) *alb4 srp54* plants, and **(B, C)** two independent lines of transgenic plants carrying the ProUBI10:GET3B-STREP transgene in an *alb4 srp54 get3b* background. Bottom row: Images of (D) *alb4 srp54 get3b* plants, and **(E, F)** two independent lines of transgenic plants carrying the ProUBI10:GET3B(D124N)-STREP transgene in an *alb4 srp54 get3b* background. Bars= 1 cm. **(G)** Rosette diameters of *alb4 srp54* plants and three independent lines of *alb4 srp54 get3b* plants carrying the ProUBI10:GET3B-STREP transgene. **(H)** Rosette diameters of untransformed (T0) *alb4 srp54 get3b* plants and four independent lines of *alb4 srp54 get3b* plants carrying the ProUBI10:GET3B(D124N)-STREP transgene. Data are presented as means + SE (n= 8-10 individual plants in G, n= 3-8 individual plants in H). Asterisks designate significant differences (Student’s t test, *P<0.05, **P<0.01) between the transgenic plants and either *alb4 srp54* plants (G) or *alb4 srp54* get3b plants (H).

To determine if the molecular complementation was due to the targeting or holdase function of GET3B, we introduced *GET3B* transgenes with D124N mutations into the *alb4 srp54 get3b* background (Figure 5D-F). These have also been previously used (Anderson et al., 2021) to eliminate the ATPase activity of GET3B (Mateja et al., 2009b). In the T2 generation 7 independent lines were identified that contained the *ProUB10:GET3B (D124N)-STREP* transgene. Data are shown for 4 randomly chosen lines. One of the lines (line2) displayed a significantly larger rosette diameter than the triple mutant but all of the lines were approximately three times smaller than the *alb4 srp54* double mutant (Figure 5D-H). Overall, the transgene with a D124N mutation was unable to rescue the *alb4 srp54 get3b* phenotype, illustrating that the ATP hydrolysis activity is crucial for complementation (Figure 5H).

To further investigate the functional consequences of the WT GET3B transgene and the ATP hydrolysis variant, we transfected these constructs into *alb4* or *stic2* backgrounds. In the T2 generation, we observed distinct phenotypic differences between the untransfected and transgenic lines (Figure 6). Transfection of the WT GET3B transgene into *stic2* backgrounds did not result in significant phenotypic differences compared to *stic2* plants (Figure 6C, D, G). In contrast, transfection of the ATP hydrolysis-deficient GET3B variant into the *alb4* background led to a significantly smaller plant size in one of two randomly chosen lines (Figure 6E, H). This effect was more pronounced when the transgene was transfected into a *stic2* background. Both of the two randomly chosen lines showed highly significant reductions in rosette diameter when compared to *stic2* plants (Figure 6F, H).

**Figure 6.**
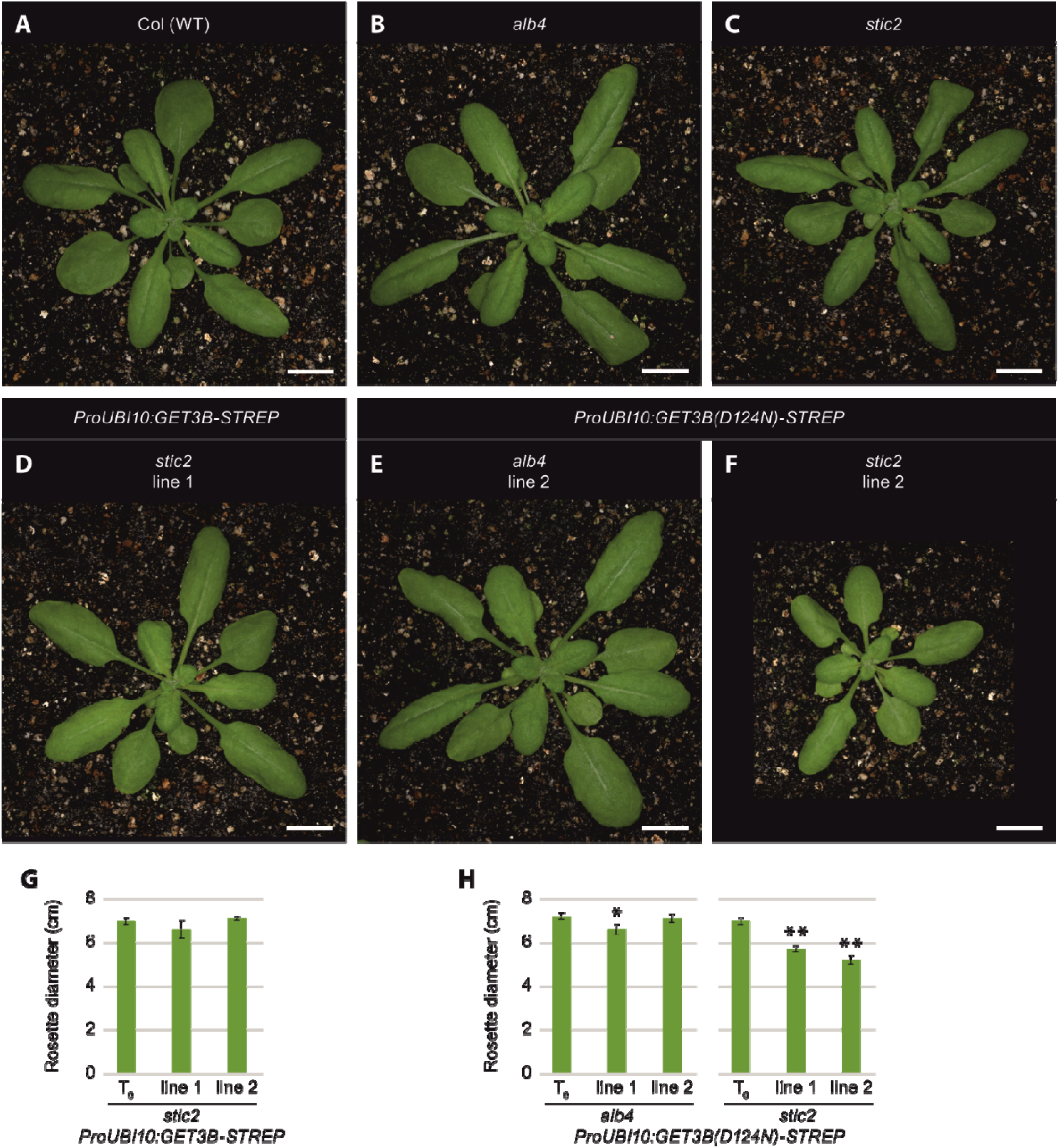
Expression of transgenes encoding GET3B with a D124N mutation leads to reductions in plant size in alb4 or stic2 genetic backgrounds. Plants were grown in 12-h light/12-h dark conditions for 4 weeks. Top row: Images of **(A)** Columbia wild-type, (B) *alb4*, and (C) *stic2* plants. Bottom row: Transgenic plants carrying the **(D)** ProUBI10:GET3B-STREP transgene in a *stic2* background, **(E)** ProUBI10:GET3B(D124N)-STREP transgene in an *alb4* background and **(F)** ProUBI10:GET3B(D124N)-STREP transgene in a *stic2* background. Bars= 1 cm. **(G)** Rosette diameters of untransformed (T0) *stic2* plants and two independent lines of transgenic *stic2* plants carrying the ProUBI10:GET3B-STREP transgene. **(H)** Rosette diameters of untransformed (T0) alb4 (left) or *stic2* (right) plants and two independent lines each of transgenic *alb4* or *stic2* plants carrying the ProUBI10:GET3B(D124N)-STREP transgene. Data are presented as means + SE (n= 10-14 individual plants in G, n= 10-15 individual plants in H). Asterisks indicate significant differences (Student’s t test, *P<0.05, **P<0.01) between the transgenic plants and untransformed (T0) plants of the same genotype.

These findings show that the loss of ATP hydrolysis of GET3B in conjunction with impairments of the STIC pathway result in stunted growth. This clearly illustrates the importance of both functioning pathways for normal plant growth and chloroplast function.

### GET3B interacts physically with the C-termini of ALB3 and ALB4

Due to the fact that the *alb3* mutant does not survive beyond the seedling stage when germinated on soil (Sundberg et al., 1997), we did not include it in our preceding genetic analysis. Still we wanted to analyze the interplay of GET3B with ALB3 as well as ALB4 and verify if the latter two could function as the terminal stromal insertases of the pathway. We were interested in both homologues since these were suggested to engage with similar sets of interactors (Trösch et al., 2015).

To investigate the interaction between GET3B and ALB3 or ALB4, we recombinantly expressed GET3B C-terminally fused to GST (GST-Get3b) as well as C-terminal fragments of ALB3 and ALB4 N-terminally fused to a His-tag (Alb3C-His and Alb4C-His; Ackermann et al., 2021) and performed pulldown experiments (Figure 7A, B). We mixed GST-Get3b with Alb3-His as well as GST-Get3b with Alb4C-His and incubated them with Glutathione Sepharose. In order to check for unspecific interactions, Alb3C- and Alb4C-His were incubated with GST and Glutathione Sepharose resin (Figure 7A). We observed residual amounts of both Alb3C- and Alb4C-His being pulled down by Glutathione Sepharose and the GST tag, indicating a non-specific interaction with the resin. However, when GST-Get3b was included in the pulldown experiment, we observed a clear enrichment of both Alb3C- and Alb4C-His, suggesting a specific interaction between GET3B and the C-terminus of ALB3 and ALB4 (Figure 7B).

**Figure 7.**
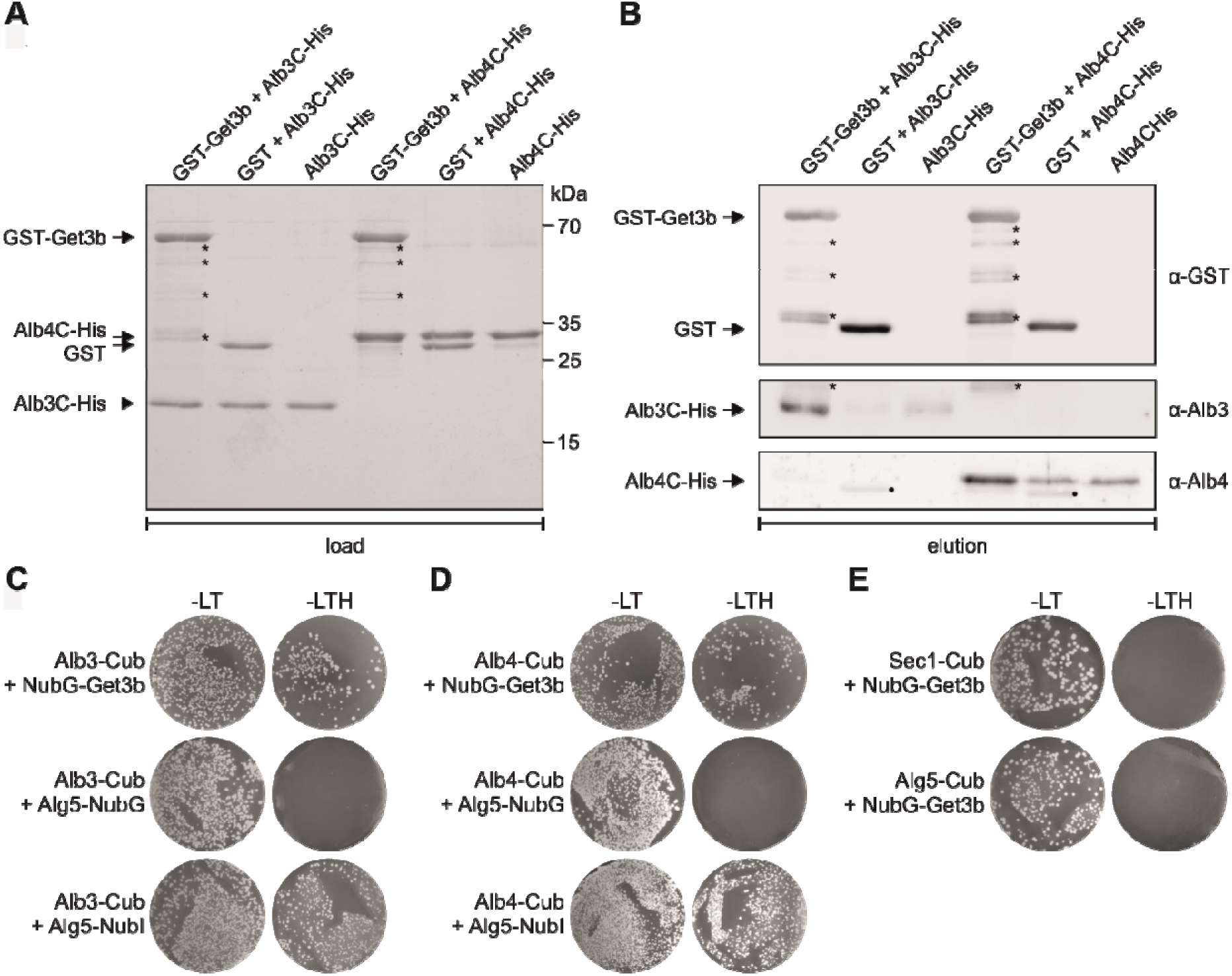
Get3b interacts with the thylakoid membrane proteins Alb3 and Alb4. **(A)** In vitro pull-down assays were performed with recombinant GST-tagged mature Get3b (GST-Get3b) and the His-tagged C-terminus of Alb3 (Alb3C-His) or Alb4 (Alb4C-His). Samples containing GST and the C-termini, or the C-termini alone served as negative controls. Approximately 4% of the load samples were applied to SDS-PAGE analysis and stained with Coomassie. **(B)** 10% of the eluates of the pull-down assay were subjected to immunoblot analysis using antibodies specific for Alb3, Alb4, or GST. The asterisks indicate truncated forms of the GST-Get3b. The closed circle indicates a nonspecific reaction of the anti-Alb4 antibody with GST. **(C)** Split-ubiquitin yeast two-hybrid assay using mature Get3b and mature full-length Alb3. Alb3 was used as fusion with the C-terminal half of ubiquitin (Cub) at its C-terminus (Alb3-Cub). Get3b was used as fusion with a modified N-terminal half of ubiquitin (NubG) at its N-terminus (NubG-Get3b). The unrelated endoplasmic reticulum membrane protein Alg5 fused to wild-type Nub (Alg5-NubI) and NubG (Alg5-NubG) served as positive and negative controls for the Cub-fusion construct, respectively. DSY-1 yeast colonies were first plated on permissive medium (−LT, lacking Leu and Trp) and on selective medium (−LTH, lacking Leu, Trp, and His). **(D)** Same assay as in (C) but using mature full-length Alb4 as bait. **(E)** Split-ubiquitin interaction tests of SecY1-Cub with NubG-Get3b and of Alg5-Cub with NubG-Get3b as negative control.

In order to corroborate the *in vitro* findings we employed a yeast two-hybrid split ubiquitin assay to further investigate the interaction between GET3B and ALB3 as well as ALB4. We transfected yeast cells with constructs expressing full length ALB3 or ALB4 N-terminally fused to the C-terminus of ubiquitin (Alb3-Cub and Alb4-Cub), along with GET3B fused to the N-terminus of ubiquitin (NubG-Get3b) (Figure 7C, D). As controls, we transfected cells with Alg5 fused to the N-terminus of ubiquitin, both with and without the mutation allowing spontaneous reassembly of ubiquitin (Alg5-NubG and Alg5-NubI, respectively). As additional controls, we tested the C-terminus of ubiquitin N-terminally fused to SEC1 and ALG5 (Sec1-Cub and Alg5-Cub) together with NubG-Get3b (Figure 7E).

Upon growth on the corresponding dropout medium, we observed robust cell growth in combination with NubG-Get3b and Alb3-Cub as well as Alb4-Cub fusion constructs, indicating a positive interaction (Figure 7C, D second column). In contrast, cells co-transfected with the control protein, Alg5-NubG as well as Sec1-Cub did not exhibit significant growth under the same conditions (Figure 7E second column). As positive controls, cells expressing Alg5-NubG grew on all media, confirming the functionality of the assay.

The combination of the pulldown experiments as well as the yeast two-hybrid split ubiquitin experiment indicated a physical interaction between GET3B and the C-terminus of ALB3 and ALB4. These results together with previous ones (Anderson et al., 2021) support the notion that GET3B selectively binds clients and interacts with insertases, potentially facilitating the membrane integration of proteins within chloroplasts.

## Discussion

The findings of this study illuminate the critical role of *At*GET3B in chloroplast function. We provide new mechanistic insights of its function during photosynthesis and chloroplast biogenesis in *Arabidopsis thaliana*.

### Impact of GET3B on chloroplast proteome, PS II assembly and carbon fixation

The proteomic analysis provided a comprehensive overview of the downstream effects caused by the absence or reduction of GET3B. This was crucial in identifying the specific proteins and pathways most affected by the *get3b* mutation, shedding light on the multifaceted role of GET3B in chloroplast biogenesis and function. One of the most striking findings from the proteomic analysis was the global effect on the abundance of plastudic proteins in the *get3b* mutant (Figure 1). These encompass a wide functional range, suggesting that GET3B has a broad impact on chloroplast proteostasis.

The entire photosynthetic apparatus of higher plats contains 16 single pass IMPs (Shi and Schröder, 2004): PS II contains 12 of which 7 have a TA topology (Wei et al., 2016) PS I comprises two of which none have a TA topology (Pan et al., 2018) and the ATP synthase encompasses two of which one shares a TA topology (Hahn et al., 2018). Taking the postulated function of GET3B into account as the central element of the stromal TA targeting machinery one would expect the corresponding substrate to be reduced in a genetic background of a knock-out or knock-down. Remarkably, the only single pass IMP that was significantly depleted in *get3b* was PsbR (Supplemental table S1). Together with PsbO, PsbP and PsbQ, PsbR forms the OEC and represents its only IMP (Mishra and Ghanotakis, 1993). To date structural analyses of PS II of higher plants were unable to resolve the location and topology of PsbR (Van Bezouwen et al., 2017). However, previous biochemical analyses corroborated that the majority of the elongated N-terminus is oriented to the thylakoidal lumen (Ljungberg et al., 1984; Ljungberg et al., 1986). If this topology is correct, then no bona fide photosynthetic TA protein was significantly depleted by the absence or reduction of GET3B. This implies that either GET3B is not involved in the targeting of TA proteins or another factor can compensate for its absence (see next section: genetic and physical interactions with STIC component and SRP54).

The significant reduction of Fv/Fm values during the de novo assembly of PS II highlight the important role of GET3B in early chloroplast biogenesis. The partial restoration of emitted fluorescence and concomitant PS II functionality in the presence of an external carbon source suggests that GET3B may play a crucial role under autotrophic conditions (Figure 2A-C). The observed reduction of Fv/Fm values during de-etiolation could either arise from impediments in OEC assembly/function or the depletion of key photosynthetic assembly factors (Supplemental table S1). It is difficult to dissect whether this reduction actually results from an incomplete assembly of PS II or solely an impeded OEC of PS II. Further analyses are necessary to clarify this.

Components of chlorophyll biogenesis pathways such as CHL synthase, magnesium chelatase and several assembly factors as well as subunits of RuBisCo were significantly depleted in *get3b* (Supplemental table S1). This directly correlates with the reduced chlorophyll content of young *get3b* seedlings (Figure 2D) as well as the reduced carbon assimilation (Figure 3), respectively. These observations further support the conclusion that GET3B is integral to the proper formation and function of the entire photosynthetic apparatus. However, at this stage it is hard to dissect if these are due to the direct function of GET3B or secondary pleiotropic effects.

The proteomic analysis provided critical insight into the molecular underpinnings of the *get3b* mutation. GET3B appears to be vital for the maintenance of a wide range of chloroplast proteins, particularly those involved in photosynthesis. The broad impact of GET3B on chloroplast proteome underscores its central role in chloroplast biogenesis and function, with potential implications for understanding how chloroplast proteostasis is maintained under various physiological conditions.

### Genetic and physical interactions with STIC components and SRP54

The negative synergism derived from combining the *get3b* allele with *srp54*, *alb4* and *stic2* implies an orchestrated network between GET3B and these pathway components probably by ensuring the proper integration and assembly of chloroplast proteins.

In particular, the *srp54 get3b* double mutant displayed a more severe phenotype than either single mutant by itself (Figure 4), signifying that the *srp54* mutation acts as a potentiator of the *get3b* effect or vice versa. This suggests that SRP54 and GET3B either may work together in a single pathway or may work separately in parallel pathways that converge in similar chloroplastidic processes. Interestingly, SRP54 binds solely to the third TMD of LHCPs (High et al., 1997). This TMD is the most hydrophobic of all three and fusing it alone to a soluble protein will translocate the fusion protein into the thylakoid membrane (Kohorn and Tobin, 1989). Since both SRP54 and GET3B are capable of binding single TMDs, it might be the case that these two proteins are able to partially compensate for the absence of the other, thereby maintaining stromal IMP trafficking when other targeting networks are not fully functional.

The incapacity of the ATPase-deficient variant to rescue the growth phenotype in the *alb4 srp54 get3b* triple mutant (Figure 5) underscores that it is the targeting function of GET3B rather than the potential holdase function which enables its participation in chloroplastidic proteostasis. The observed growth defects of plants expressing the ATPase-defiant variant in *alb4* and *stic2* backgrounds (Figure 6) further confirm that operative ATPase activity is essential for a fully functioning GET3B in planta.

The physical interaction between GET3B and the C-terminal regions of ALB3 and ALB4, as demonstrated by pulldown experiments and yeast two-hybrid assays, provides a mechanistic insight into how GET3B may facilitate the insertion of proteins into the thylakoid membrane. Interestingly, irrespective of *in vitro* or *in vivo,* both the stromal exposed C-terminal regions or full length ALB3 and ALB4 interacted with GET3B to a fairly similar degree (Figure 7), indicating less selectivity for the insertases than the cpSRP pathway. The C-terminal regions of ALB3 and ALB4 share two conserved motifs: motif I and motif III, which are not utilized by ALB3 for the interaction with SRP43 (Falk et al., 2010). In turn, SRP43 interacts with motifs II and IV of ALB3 which helps it discriminate between the two insertases (Falk et al., 2010; Dünschede et al., 2011). Recently, a direct physical interaction between STIC2 and motif III of ALB3 and ALB4 was demonstrated (Stolle et al., 2024). The plastidic ribosome on the other hand cooperatively binds motif III and IV of ALB3 (Ackermann et al., 2021). In future studies, it would be interesting to characterize the GET3B binding site within the C-terminal regions of the Alb proteins and to investigate whether GET3B and STIC2 may engage in competitive or cooperative binding.

In summary, the results of this study highlight the role of GET3B in chloroplast biogenesis and function. GET3B appears to be involved in the proper assembly and function of photosynthetic complexes but also for proteostasis in a broader sense within the chloroplast. Its genetic interactions with STIC components and SRP54, as well as its dependence on ATPase activity, suggests a critical role in the integration of membrane proteins. Further studies are necessary to explore the precise mechanistic details of GET3B**’**s interactions with the other targeting components and how they contribute to the overall regulation of chloroplast biogenesis and function.

## Acknowledgements

The authors would like thank Sarah Friedrich for plant photographs and figure preparation as well as Fabian Fuchs and Silke Funke for experimental support.

## Author contribution

Uwe Bodensohn, Danja Schünemann, Donna Fernandez and Enrico Schleiff conceived the idea and designed the research. Donna Fernandez executed the genetic analysis. Beatrix Dünschede and Chiara Kuhlmann performed the *in vivo* and *in vitro* interaction investigations. Khushbu Kumari implemented the IRGA measurements. Roman Ladig performed the LC-MS/MS analysis. Uwe Bodensohn executed the physiological experiments and drafted the manuscript. All authors commented on the previous versions of the manuscript. All authors approved the final manuscript.

## Data availability

The datasets generated during and/or analyzed during the current study are available from the corresponding author on reasonable request.

## References

Ackermann, B., Dünschede, B., Pietzenuk, B., Justesen, B.H., Krämer, U., Hofmann, E., Günther Pomorski, T., and Schünemann, D. (2021). Chloroplast Ribosomes Interact With the Insertase Alb3 in the Thylakoid Membrane. Front. Plant Sci. 12.

Aigner, H., Wilson, R.H., Bracher, A., Calisse, L., Bhat, J.Y., Hartl, F.U., and Hayer-Hartl, M. (2017). Plant RuBisCo assembly in E. coli with five chloroplast chaperones including BSD2. Science (80-. ). 358: 1272–1278.

Allahverdiyeva, Y. et al. (2013). Arabidopsis plants lacking PsbQ and PsbR subunits of the oxygen-evolving complex show altered PSII super-complex organization and short-term adaptive mechanisms. Plant J. 75: 671–684.

Amin, P., Sy, D.A.C., Pilgrim, M.L., Parry, D.H., Nussaume, L., and Hoffman, N.E. (1999). Arabidopsis mutants lacking the 43- and 54-kilodalton subunits of the chloroplast signal recognition particle have distinct phenotypes. Plant Physiol. 121: 61–70.

Anderson, S.A., Satyanarayan, M.B., Wessendorf, R.L., Lu, Y., and Fernandez, D.E. (2021). A homolog of GuidedEntry of Tail-anchored proteins3 functions in membrane-specific protein targeting in chloroplasts of Arabidopsis. Plant Cell 33: 2812–2833.

Anghel, S.A., McGilvray, P.T., Hegde, R.S., and Keenan, R.J. (2017). Identification of Oxa1 Homologs Operating in the Eukaryotic Endoplasmic Reticulum. Cell Rep. 21: 3708–3716.

Bals, T., Dünschede, B., Funke, S., and Schünemann, D. (2010). Interplay between the cpSRP pathway components, the substrate LHCP and the translocase Alb3: An in vivo and in vitro study. FEBS Lett. 584: 4138–4144.

Barlow, A.N., Manu, M.S., Saladi, S.M., Tarr, P.T., Yadav, Y., Thinn, A.M.M., Zhu, Y., Laganowsky, A.D., Clemons, W.M., and Ramasamy, S. (2023). Structures of Get3d reveal a distinct architecture associated with the emergence of photosynthesis. J. Biol. Chem. 299: 104752.

Bauer, J., Chen, K., Hiltbunner, A., Wehrli, E., Eugster, M., Schnell, D., and Kessler, F. (2000). The major protein import receptor of plastids is essential for chloroplast biogenesis. Nature 403: 203–207.

Bédard, J., Trösch, R., Wu, F., Ling, Q., Flores-Pérez, Ú., Töpel, M., Nawaz, F., and Jarvis, P. (2017). Suppressors of the chloroplast protein import mutant tic40 reveal a genetic link between protein import and thylakoid biogenesis. Plant Cell 29: 1726–1747.

Benz, M., Bals, T., Gügel, I.L., Piotrowski, M., Kuhn, A., Schünemann, D., Soll, J., and Ankele, E. (2009). Alb4 of arabidopsis promotes assembly and stabilization of a non chlorophyll-binding photosynthetic complex, the CF1CF0-ATP synthase. Mol. Plant 2: 1410–1424.

van Berkel, A.A., Santos, T.C., Shaweis, H., van Weering, J.R.T., Toonen, R.F., and Verhage, M. (2020). Loss of MUNC18-1 leads to retrograde transport defects in neurons. J. Neurochem.

Van Bezouwen, L.S., Caffarri, S., Kale, R., Kouřil, R., Thunnissen, A.M.W.H., Oostergetel, G.T., and Boekema, E.J. (2017). Subunit and chlorophyll organization of the plant photosystem II supercomplex. Nat. Plants 3: 1–11.

Bodensohn, U.S., Simm, S., Fischer, K., Jäschke, M., Groß, L.E., Kramer, K., Ehmann, C. stian, Rensing, S.A., Ladig, R., and Schleiff, E. (2019). The intracellular distribution of the components of the GET system in vascular plants. Biochim. Biophys. Acta - Mol. Cell Res. 1866: 1650–1662.

Bozkurt, G., Stjepanovic, G., Vilardi, F., Amlacher, S., Wild, K., Bange, G., Favaloro, V., Rippe, K., Hurt, E., Dobberstein, B., and Sinning, I. (2009). Structural insights into tail-anchored protein binding and membrane insertion by Get3. Proc. Natl. Acad. Sci. U. S. A. 106: 21131–21136.

Brettel, K. (1997). Electron transfer and arrangement of the redox cofactors in photosystem I. Biochim. Biophys. Acta - Bioenerg. 1318: 322–373.

Cox, J. and Mann, M. (2008). MaxQuant enables high peptide identification rates, individualized p.p.b.-range mass accuracies and proteome-wide protein quantification. Nat. Biotechnol. 26: 1367–1372.

Duncan, O., van der Merwe, M.J., Daley, D.O., and Whelan, J. (2013). The outer mitochondrial membrane in higher plants. Trends Plant Sci. 18: 207–217.

Dünschede, B., Bals, T., Funke, S., and Schünemann, D. (2011). Interaction studies between the chloroplast signal recognition particle subunit cpSRP43 and the full-length translocase Alb3 reveal a membrane-embedded binding region in Alb3 protein. J. Biol. Chem. 286: 35187–35195.

Falk, S., Ravaud, S., Koch, J., and Sinning, I. (2010). The C terminus of the Alb3 membrane insertase recruits cpSRP43 to the thylakoid membrane. J. Biol. Chem. 285: 5954–5962.

Gerdes, L., Bals, T., Klostermann, E., Karl, M., Philippar, K., Hünken, M., Soll, J., and Schünemann, D. (2006). A second thylakoid membrane-localized Alb3/OxaI/YidC homologue is involved in proper chloroplast biogenesis in Arabidopsis thaliana. J. Biol. Chem. 281: 16632–16642.

Hackett, J.B., Shi, X., Kobylarz, A.T., Lucas, M.K., Wessendorf, R.L., Hines, K.M., Bentolila, S., Hanson, M.R., and Lu, Y. (2017). An organelle RNA recognition motif protein is required for photosystem II subunit psbF transcript editing. Plant Physiol. 173: 2278–2293.

Hahn, A., Vonck, J., Mills, D.J., Meier, T., and Kühlbrandt, W. (2018). Structure, mechanism, and regulation of the chloroplast ATP synthase. Science 360.

High, S., Henry, R., Mould, R.M., Valent, Q., Meacock, S., Cline, K., Gray, J.C., and Luirink, J. (1997). Chloroplast SRP54 interacts with a specific subset of thylakoid precursor proteins. J. Biol. Chem. 272: 11622–11628.

Hoagland, D. and Arnon, D. (1950). The water-culture method for growing plants without soil. Circ. Calif. Agric. Exp. Stn. 347.

Hristou, A., Gerlach, I., Stolle, D.S., Neumann, J., Bischoff, A., Dünschede, B., Nowaczyk, M.M., Zoschke, R., and Schünemann, D. (2019). Ribosome-Associated Chloroplast SRP54 Enables Efficient Cotranslational Membrane Insertion of Key Photosynthetic Proteins. Plant Cell 31: 2734–2750.

Jarvis, P. and López-Juez, E. (2013). Biogenesis and homeostasis of chloroplasts and other plastids. Nat. Rev. Mol. Cell Biol. 14: 787–802.

Jiang, X., Jiang, H., Shen, Z., and Wang, X. (2014). Activation of mitochondrial protease OMA1 by Bax and Bak promotes cytochrome c release during apoptosis. Proc. Natl. Acad. Sci. U. S. A. 111: 14782–14787.

Kalbfleisch, T., Cambon, A., and Wattenberg, B.W. (2007). A bioinformatics approach to identifying tail-anchored proteins in the human genome. Traffic 8: 1687–1694.

Kohorn, B.D. and Tobin, E.M. (1989). A hydrophobic, carboxy-proximal region of a light-harvesting chlorophyll a/b protein is necessary for stable integration into thylakoid membranes. Plant Cell 1: 159–166.

León, I.R., Schwämmle, V., Jensen, O.N., and Sprenger, R.R. (2013). Quantitative assessment of in-solution digestion efficiency identifies optimal protocols for unbiased protein analysis. Mol. Cell. Proteomics 12: 2992–3005.

Link, A.J., Skretas, G., Strauch, E.-M., Chari, N.S., and Georgiou, G. (2008). Efficient production of membrane-integrated and detergent-soluble G protein-coupled receptors in Escherichia coli. Protein Sci. 17: 1857–1863.

Liu, H., Li, A., Rochaix, J.D., and Liu, Z. (2023). Architecture of chloroplast TOC–TIC translocon supercomplex. Nature 615: 349–357.

Ljungberg, U., Åkerlund, H. -E, and Andersson, B. (1986). Isolation and characterization of the 10-kDa and 22-kDa polypeptides of higher plant photosystem 2. Eur. J. Biochem. 158: 477–482.

Ljungberg, U., Åkerlund, H.E., and Andersson, B. (1984). The release of a 10-kDa polypeptide from everted photosystem II thylakoid membranes by alkaline tris. FEBS Lett. 175: 255–258.

Mateja, A., Szlachcic, A., Downing, M.E., Dobosz, M., Mariappan, M., Hegde, R.S., and Keenan, R.J. (2009a). The structural basis of tail-anchored membrane protein recognition by Get3. Nature 461: 361–366.

Mateja, A., Szlachcic, A., Downing, M.E., Dobosz, M., Mariappan, M., Hegde, R.S., and Keenan, R.J. (2009b). The structural basis of tail-anchored membrane protein recognition by Get3. Nature 461: 361–366.

Mishra, R.K. and Ghanotakis, D.F. (1993). Selective extraction of 22 kDa and 10 kDa polypeptides from Photosystem II without removal of 23 kDa and 17 kDa extrinsic proteins. Photosynth. Res. 36: 11–16.

Moore, M., Goforth, R.L., Mori, H., and Henry, R. (2003). Functional interaction of chloroplast SRP/FtsY with the ALB3 translocase in thylakoids: Substrate not required. J. Cell Biol. 162: 1245–1254.

Mukhopadhyay, R., Ho, Y.S., Swiatek, P.J., Rosen, B.P., and Bhattacharjee, H. (2006). Targeted disruption of the mouse Asna1 gene results in embryonic lethality. FEBS Lett. 580: 3889–3894.

Murashige, T. and Skoog, F. (1962). A Revised Medium for Rapid Growth and Bio Assays with Tobacco Tissue Cultures. Physiol. Plant. 15: 473–497.

Norris, S.R., Meyer, S.E., and Callis, J. (1993). The intron of Arabidopsis thaliana polyubiquitin genes is conserved in location and is a quantitative determinant of chimeric gene expression. Plant Mol. Biol. 21: 895–906.

Pan, X., Ma, J., Su, X., Cao, P., Chang, W., Liu, Z., Zhang, X., and Li, M. (2018). Structure of the maize photosystem I supercomplex with light-harvesting complexes I and II. Science (80-. ). 360: 1109–1113.

Pasch, J.C., Nickelsen, J., and Schünemann, D. (2005). The yeast split-ubiquitin system to study chloroplast membrane protein interactions. Appl. Microbiol. Biotechnol. 69: 440–447.

Pilgrim, M.L., Van Wijk, K.J., Parry, D.H., Sy, D.A.C., and Hoffman, N.E. (1998). Expression of a dominant negative form of cpSRP54 inhibits chloroplast biogenesis in Arabidopsis. Plant J. 13: 177–186.

Powis, K., Schrul, B., Tienson, H., Gostimskaya, I., Breker, M., High, S., Schuldiner, M., Jakob, U., and Schwappach, B. (2013). Get3 is a holdase chaperone and moves to deposition sites for aggregated proteins when membrane targeting is blocked. J. Cell Sci. 126: 473–483.

Rappsilber, J., Mann, M., and Ishihama, Y. (2007). Protocol for micro-purification, enrichment, pre-fractionation and storage of peptides for proteomics using StageTips. Nat. Protoc. 2: 1896–1906.

Schindelin, J. et al. (2012). Fiji: an open-source platform for biological-image analysis. Nat. Methods 9: 676–682.

Schuenemann, D., Gupta, S., Persello-Cartieauxi, F., Klimyuk, V.I., Jones, J.D.G., Nussaume, L., and Hoffman, N.E. (1998). A novel signal recognition particle targets light-harvesting proteins to the thylakoid membranes. Proc. Natl. Acad. Sci. U. S. A. 95: 10312–10316.

Schuldiner, M., Metz, J., Schmid, V., Denic, V., Rakwalska, M., Schmitt, H.D., Schwappach, B., and Weissman, J.S. (2008). The GET Complex Mediates Insertion of Tail-Anchored Proteins into the ER Membrane. Cell 134: 634–645.

Shi, L.X. and Schröder, W.P. (2004). The low molecular mass subunits of the photosynthetic supracomplex, photosystem II. Biochim. Biophys. Acta - Bioenerg. 1608: 75–96.

Sommer, M., Rudolf, M., Tillmann, B., Tripp, J., Sommer, M.S., and Schleiff, E. (2013). Toc33 and Toc64-III cooperate in precursor protein import into the chloroplasts of Arabidopsis thaliana. Plant, Cell Environ. 36: 970–983.

Stolle, D.S. et al. (2024). STIC2 selectively binds ribosome-nascent chain complexes in the cotranslational sorting of Arabidopsis thylakoid proteins. EMBO J.

Sundberg, E., Slagter, J.G., Fridborg, I., Cleary, S.P., Robinson, C., and Coupland, G. (1997). ALBINO3, an arabidopsis nuclear gene essential for chloroplast differentiation, encodes a chloroplast protein that shows homology to proteins present in bacterial membranes and yeast mitochondria. Plant Cell 9: 717–730.

Toro, G., Flexas, J., and Escalona, J.M. (2019). Contrasting leaf porometer and infra-red gas analyser methodologies: an old paradigm about the stomatal conductance measurement. Theor. Exp. Plant Physiol. 31: 483–492.

Trösch, R., Töpel, M., Flores-Pérez, Ú., and Jarvis, P. (2015). Genetic and physical interaction studies reveal functional similarities between ALBINO3 and ALBINO4 in Arabidopsis. Plant Physiol. 169: 1292–1306.

Tseng, Y.Y., Yu, C.W., and Liao, V.H.C. (2007). Caenorhabditis elegans expresses a functional ArsA. FEBS J. 274: 2566–2572.

Tyanova, S. and Cox, J. (2018). Perseus: A Bioinformatics Platform for Integrative Analysis of Proteomics Data in Cancer Research. Methods Mol. Biol. 1711: 133–148.

Vermeulen, M., Hubner, N.C., and Mann, M. (2008). High confidence determination of specific protein-protein interactions using quantitative mass spectrometry. Curr. Opin. Biotechnol. 19: 331–337.

Voth, W., Schick, M., Gates, S., Li, S., Vilardi, F., Gostimskaya, I., Southworth, D.R., Schwappach, B., and Jakob, U. (2014). The protein targeting factor Get3 functions as ATP-Independent chaperone under oxidative stress conditions. Mol. Cell 56: 116–127.

Wallin, E. and Von Heijne, G. (1998). Genome-wide analysis of integral membrane proteins from eubacterial, archaean, and eukaryotic organisms. Protein Sci. 7: 1029–1038.

Wang, F., Brown, E.C., Mak, G., Zhuang, J., and Denic, V. (2010). A chaperone cascade sorts proteins for posttranslational membrane insertion into the endoplasmic reticulum. Mol. Cell 40: 159–171.

Wang, F., Chan, C., Weir, N.R., and Denic, V. (2014). The Get1/2 transmembrane complex is an endoplasmic-reticulum membrane protein insertase. Nature 512: 441–444.

Wei, X., Su, X., Cao, P., Liu, X., Chang, W., Li, M., Zhang, X., and Liu, Z. (2016). Structure of spinach photosystem II-LHCII supercomplex at 3.2 Å resolution. Nature 534: 69–74.

Wellburn, A.R. (1994). The spectral determination of chlorophylls a and b, as well as total carotenoids, using various solvents with spectrophotometers of different resolution. J. Plant Physiol. 144: 307–313.

Xing, S., Mehlhorn, D.G., Wallmeroth, N., Asseck, L.Y., Kar, R., Voss, A., Denninger, P., Schmidt, V.A.F., Schwarzländer, M., Stierhof, Y.D., Grossmann, G., and Grefen, C. (2017). Loss of GET pathway orthologs in Arabidopsis thaliana causes root hair growth defects and affects SNARE abundance. Proc. Natl. Acad. Sci. U. S. A. 114: E1544–E1553.

Yu, B., Gruber, M., Khachatourians, G.G., Zhou, R., Epp, D.J., Hegedus, D.D., Parkin, I.A.P., Welsch, R., and Hannoufa, A. (2012). Arabidopsis cpSRP54 regulates carotenoid accumulation in Arabidopsis and Brassica napus. J. Exp. Bot. 63: 5189–5202.

Ziehe, D., Dünschede, B., and Schünemann, D. (2018). Molecular mechanism of SRP-dependent light-harvesting protein transport to the thylakoid membrane in plants. Photosynth. Res. 138: 303–313.

