## Supplemental File for "GET3B interacts with the thylakoidal ALB3 and ALB4 insertases and is involved in the initial stages of chloroplast biogenesis"

**Supplemental data**

**Supplemental tables**

**Table S1. Listed significant proteinaceous hits from the comparative proteomics.** These are grouped according to their cellular function and depleted as well as enriched hits are marked with 🡫 or 🡩, respectively. This table supports Fig. 1 of the manuscript and significantly contributes to the transparency of the identified proteinaceous hits of the paper.

| Accession | ID | Cellular function | Abundance | |
| --- | --- | --- | --- | --- |
| At3g01510 | Putative phosphatase (AtLSF1) | Other | 🡫 |  |
| At3g56130 | Biotin/lipoyl attachment domain protein (AtBADC1) | Other | 🡫 |  |
| At1g12900 | Glyceraldehyde 3-phosphate dehydrogenase A subunit (AtGAPa-2) | Other | 🡫 |  |
| At1g31860 | Encodes a bifunctional protein (AtHISN2) | Other | 🡫 |  |
| At1g31180 | Encodes a isopropylmalate dehydrogenase (AtIMD3) | Other | 🡫 |  |
| At5g36870 | Encodes a gene similar to callose synthase (AtCalS4) | Other | 🡫 |  |
| At5g57850 | Encodes a involved in D-Amino Acid production (AtADCL) | Other | 🡫 |  |
| At3g49680 | Encodes branched-chain amino acid aminotransferase (AtBCAT-3) | Other | 🡫 |  |
| At1g48860 | 5-enolpyruvylshikimate-3-phosphate synthase (EPSP) | Other | 🡫 |  |
| At4g16155 | Dihydrolipoamide dehydrogenase (AtptLPD2) | Other | 🡫 |  |
| At4g31990 | Encodes a plastid-localized aspartate aminotransferase (AtASP5) | Other | 🡫 |  |
| At5g04740 | Encodes a ACT domain-containing protein (AtACR12) | Other | 🡫 |  |
| At5g60600 | Encodes a chloroplast-localized HDS protein (AtHDS) | Other | 🡫 |  |
| At2g47400 | GAPDH regulatory protein (AtCP12-1) | Other | 🡫 |  |
| At4g15530 | Dual-targeted protein pyruvate, orthophosphate dikinase.(AtPPDK) | Other | 🡫 |  |
| At1g01090 | Pyruvate dehydrogenase E1 alpha subunit (AtPDH-E1) | Other | 🡫 |  |
| At1g75330 | Ornithine carbamoyltransferase (AtOTC) | Other | 🡫 |  |
| At5g63310 | Nucleoside diphosphate kinase (AtNDPK2) | Other | 🡫 |  |
| At4g00620 | Amino acid dehydrogenase protein (AtFOLD4) | Other | 🡫 |  |
| At2g37660 | Putative NAD-dependent epimerase/dehydratase | Other | 🡫 |  |
| At2g26930 | 4-(cytidine 5'-phospho)-2-C-methyl-D-erithritol kinase (AtCMK) | Other | 🡫 |  |
| At4g19710 | Aspartate kinase/homoserine dehydrogenase (AtAK-HsDH2) | Other | 🡫 |  |
| At2g29630 | Encodes a protein involved in thiamine biosynthesis (AtTHIC) | Other | 🡫 |  |
| At5g54810 | [Putative beta subunit of tryptophan synthase (AtTSB1)](javascript:openRefGene('63851');markCell('0');) | Other | 🡫 |  |
| At3g48560 | Catalyzes the formation of acetolactate from pyruvate (AtALS) | Other | 🡫 |  |
| At1g32060 | Phosphoribulokinase.(AtPRK) | Other | 🡫 |  |
| At3g54050 | Putative plastidial fructose-1,6-bisphosphatase (AtHCEF1) | Other | 🡫 |  |
| At3g26650 | Glyceraldehyde-3-phosphate dehydrogenase (AtGAPa-1) | Other | 🡫 |  |
| At1g42970 | Glyceraldehyde-3-phosphate dehydrogenase (AtGAPb) | Other | 🡫 |  |
| At3g01500 | Encodes a putative beta-carbonic anhydrase (AtBCA1) | Other | 🡫 |  |
| At2g29690 | Encode a functional anthranilate synthase protein (AtASA2) | Other | 🡫 |  |
| At5g63570 | Putative glutamate-1-semialdehyde aminotransferase (AtGSA1) | Other | 🡫 |  |
| At3g55800 | Sedoheptulose-1,7-bisphosphatase (SBPase) | Other | 🡫 |  |
| At4g23100 | Glutamate-cysteine ligase (AtCAD2) | Other | 🡫 |  |
| At2g43750 | [Plastidial O-acetyl-L-serine sulfohydrolase (AtOAS)](javascript:openRefGene('66424');markCell('0');) | Other | 🡫 |  |
| At1g09830 | [Phosphoribosylamine-glycine ligase (AtGARS)](javascript:openRefGene('64883');markCell('0');) | Other | 🡫 |  |
| At3g01120 | Encodes a cystathionine gamma-synthase (AtCGS1) | Other | 🡫 |  |
| AtCg00500 | Subunit of the Acetyl-CoA carboxylase complex (AtACCD) | Other | 🡫 |  |
| At1g48850 | Putative chorismate synthase | Other | 🡫 |  |
| At4g21990 | Adenylylphosphosulfate reductase & thioredoxin (AtAPR3) | Other | 🡫 |  |
| At5g17990 | [Putative phosphoribosylanthranilate transferase (AtPAT1/AtTRP1)](javascript:openRefGene('68166');markCell('0');) | Other | 🡫 |  |
| At3g58610 | Putative ketol-acid reductoisomerase | Other | 🡫 |  |
| At5g54770 | Putative thiamine thiazole synthase (AtTHI1) | Other | 🡫 |  |
| At3g52380 | Encodes a chloroplast RNA-binding protein (AtCP33A) | Other | 🡫 |  |
| At2g15620 | Involved in the second step of nitrate assimilation(AtNIR1) | Other | 🡫 |  |
| At3g54640 | Conversion of indole-3-glycerolphosphate to indole (ATTSA1) | Other | 🡫 |  |
| At5g35630 | Chloroplastic glutamine synthetase (AtGSL1) | Other | 🡫 |  |
| At1g22940 | Encodes a bifunctional enzyme (AtTH1) | Other | 🡫 |  |
| At5g10460 | Flowering time regulator (AtHDF1) | Other | 🡫 |  |
| At5g26570 | Chloroplastidic phosphoglucan, water dikinase (AtOK1) | Other | 🡫 |  |
| At1g09795 | ATP phosphoribosyl transferase (AtATP-PRT2) | Other | 🡫 |  |
| At4g20960 | [Pyrimidine deaminase (AtPyrD)](javascript:openRefGene('57081');markCell('0');) | Other | 🡫 |  |
| At1g31160 | Nucleoside 5'-phosphoramidate hydrolase (AtHINT2) | Other | 🡫 |  |
| At3g03710 | Putative polyribonucleotide nucleotidyltransferase (AtPNP1) | Other | 🡫 |  |
| At2g21370 | Putative xylulose kinase with no such activity | Other | 🡫 |  |
| At2g17265 | Encodes a homoserine kinase (HSK) | Other | 🡫 |  |
| At3g03890 | [Beta-barrel heme oxygenase, HOMOLOG OF HugZ (AtHOZ)](javascript:openRefGene('53649');markCell('0');) | Other | 🡫 |  |
| At5g16710 | [Putative dehydroascorbate reductase (AtDHAR3)](javascript:openRefGene('58763');markCell('0');) | Other | 🡫 |  |
| At2g30200 | Dual-localized malonyl-CoA:ACP transacylase (AtMCAT) | Other | 🡫 |  |
| At4g18440 | Putative adenylosuccinate lyase | Other | 🡫 |  |
| At1g14810 | [Putative aspartate semialdehyde dehydrogenase](javascript:openRefGene('64952');markCell('0');) | Other | 🡫 |  |
| At1g01080 | Putative chloroplast ribonucleoprotein (AtCP28B) | Other | 🡫 |  |
| At1g67280 | [Putative type-I nickel-dependent glyoxalase (AtGLYI6)](javascript:openRefGene('15782');markCell('0');) | Other | 🡫 |  |
| At4g33500 | Putative PP2C-type protein phosphatase (AtPP2C62) | Other | 🡫 |  |
| At4g33680 | Putative diaminopimelate aminotransferase (AtDAP) | Other | 🡫 |  |
| At4g33470 | Member of the histone deacetylase family proteins (AtHDA14) | Other | 🡫 |  |
| At4g38970 | [Putative fructose-bisphosphate aldolase (AtFBA2)](javascript:openRefGene('67890');markCell('0');) | Other | 🡫 |  |
| At4g13430 | [Large subunit of isopropylmalate isomerase dimer (AtIPMI-LSU1)](javascript:openRefGene('56702');markCell('0');) | Other | 🡫 |  |
| At5g19855 | RuBisCo complex assembly chaperone (AtRbcX) | Chaperone/assembly | 🡫 |  |
| At2g28000 | Chaperonin-60 alpha (RBCS assembly, AtCPN60A1) | Chaperone/assembly | 🡫 |  |
| At2g39730 | Rubisco activase (AtRCA) | Chaperone/assembly | 🡫 |  |
| At1g55490 | [Component of plastidial Cpn60 chaperonin complex (AtCPN60B1)](javascript:openRefGene('15307');markCell('0');) | Chaperone/assembly | 🡫 |  |
| At2g44650 | Encodes a chloroplast-localized chaperonin 10 (AtCPN10-2) | Chaperone/assembly | 🡫 |  |
| AtCg00360 | [putative photosystem I (PS-I) assembly factor (AtYCF3)](javascript:openRefGene('69213');markCell('0');) | Chaperone/assembly | 🡫 |  |
| At2g21385 | [Molecular chaperone involved in ATP synthase assembly (AtBFA3)](javascript:openRefGene('17174');markCell('0');) | Chaperone/assembly | 🡫 |  |
| At3g29185 | putative chloroplast ATP synthase assembly factor (AtBFA1) | Chaperone/assembly | 🡫 |  |
| At4g37200 | [Thioredoxin-like protein - cytochrome b_6_f assembly (AtHCF164)](javascript:openRefGene('67864');markCell('0');) | Chaperone/assembly | 🡫 |  |
| At1g22700 | PS I assembly factor (AtPYG7) | Chaperone/assembly | 🡫 |  |
| At2g34860 | DnaJ-like zinc finger protein regulating PSI assembly (AtPSA2) | Chaperone/assembly | 🡫 |  |
| At3g62030 | Nuclear-encoded chloroplast stromal cyclophilin (AtCYP20-3) | Chaperone/assembly | 🡫 |  |
| At4g03520 | Encodes a redox activated co-chaperone (AtTRX-M2) | Chaperone/assembly | 🡫 |  |
| At4g34730 | [Auxiliary factor involved in plastidial 16S rRNA maturation (AtRBF1)](javascript:openRefGene('57743');markCell('0');) | Chaperone/assembly | 🡫 |  |
| At2g47910 | [Assembly factor or stabilization of NDH complex (AtCRR6)](javascript:openRefGene('53475');markCell('0');) | Chaperone/assembly | 🡫 |  |
| At3g24430 | Assembly factor for Fe-S cluster (AtHCF101) | Chaperone/assembly | 🡫 |  |
| At5g55220 | Plastidial co-translational nascent polypeptide chaperone (AtTIG) | Chaperone/assembly | 🡫 |  |
| At1g12800 | [putative plastidial RNA chaperone (AtSDP)](javascript:openRefGene('64924');markCell('0');) | Chaperone/assembly | 🡫 |  |
| At3g45050 | DnaJ-likeTransmembrane protein (DnaJ-like) (AtDnaJE1.18) | Chaperone/assembly | 🡫 |  |
| At4g13670 | Component of plastidial PEP complex (AtDnaJE1.2) | Chaperone/assembly | 🡫 |  |
| At4g26555 | FKBP-type protein folding catalyst (AtFKBP16-1) | Chaperone/assembly | 🡫 |  |
| At5g66570 | Oxygen evolving complex subunit33 (AtOEC33/AtPsbO1) | Light reaction | 🡫 |  |
| At1g06680 | Oxygen evolving complex 23 (AtOE23/AtPsbP1) | Light reaction | 🡫 |  |
| At4g05180 | Oxygen evolving complex 17 (AtOE17/AtPsbQ2) | Light reaction | 🡫 |  |
| At1g79040 | Encodes for the 10 kDa subunit of PSII (AtPsbR) | Light reaction | 🡫 |  |
| At3g21055 | Encodes photosystem II 5 kD protein subunit (AtPsbTn) | Light reaction | 🡫 |  |
| At5g64040 | PSI subunit located entirely in the thylakoid lumen (AtPsaN) | Light reaction | 🡫 |  |
| At2g26540 | Encodes a uroporphyrinogen-III (AtUROS) | Light reaction | 🡫 |  |
| At1g03630 | Protochlorophyllide oxidoreductase protein (AtPorC) | Light reaction | 🡫 |  |
| At1g04620 | Encodes a 7-hydroxymethyl chlorophyll a reductase (AtHCAR) | Light reaction | 🡫 |  |
| At3g51820 | Chlorophyll synthetase (AtG4/AtChlG) | Light reaction | 🡫 |  |
| At4g18480 | [Magnesium-protoporphyrin-IX chelatase complex (AtCHLI-1)](javascript:openRefGene('56955');markCell('0');) | Light reaction | 🡫 |  |
| At5g45930 | Subunit of magnesium chelatase (AtCHLI-2) | Light reaction | 🡫 |  |
| At1g64680 | Encodes a carotenoid isomerase (AtD27L1) | Light reaction | 🡫 |  |
| At4g28025 | putative LHCP-related protein involved in photoprotection (AtSEP5) | Light reaction | 🡫 |  |
| At4g36810 | subunit of geranyl diphosphate synthase complex (AtGGPS11) | Light reaction | 🡫 |  |
| At1g20340 | One of two Arabidopsis plastocyanin genes (AtPetE2) | Light reaction | 🡫 |  |
| At4g27800 | [PP2C-type LHCII component phosphatase (AtTAP38)](javascript:openRefGene('67732');markCell('0');) | Light reaction | 🡫 |  |
| AtCg00140 | ATPase III subunit (AtcpATP-H) | Light reaction | 🡫 |  |
| At4g38460 | Subunit of geranyl(geranyl) diphosphate synthase (AtSSU) | Light reaction | 🡫 |  |
| At1g67700 | [Protein involved in photosystem-II photodamage repair (AtHHL1)](javascript:openRefGene('65603');markCell('0');) | Light reaction | 🡫 |  |
| At5g17870 | Plastid-specific ribosomal protein 6 precursor – like (AtPSRP6) | Ribosomes | 🡫 |  |
| At3g12080 | Encodes a putative plastid-targeted GTP-binding protein (AtEngA) | Ribosomes | 🡫 |  |
| At4g01310 | Ribosomal L5P family protein (AtcpRPL5) | Ribosomes | 🡫 |  |
| At1g05190 | Encodes the plastid 50S ribosomal protein L6 (AtcpRPL6) | Ribosomes | 🡫 |  |
| At1g07320 | Encodes a plastid ribosomal protein L4 (AtcpRPL4) | Ribosomes | 🡫 |  |
| At1g79850 | 30S chloroplast ribosomal protein S17 (AtcpRPS17) | Ribosomes | 🡫 |  |
| At3g25920 | Encodes a plastid ribosomal protein CL15 (AtcpRPL15) | Ribosomes | 🡫 |  |
| At3g27850 | 50S ribosomal protein L12-C | Ribosomes | 🡫 |  |
| AtCg00780 | Encodes a chloroplast ribosomal protein L14 (AtcpRPL14) | Ribosomes | 🡫 |  |
| AtCg00790 | Chloroplast gene encoding a ribosomal protein L16 (AtcpRPL16) | Ribosomes | 🡫 |  |
| AtCg00810 | Encodes a chloroplast ribosomal protein L22 (AtcpRPL22) | Ribosomes | 🡫 |  |
| AtCg00640 | Encodes a chloroplast ribosomal protein L33 (AtcpRPL33) | Ribosomes | 🡫 |  |
| AtCg00160 | Chloroplast ribosomal protein S2 (AtcpRPS2) | Ribosomes | 🡫 |  |
| AtCg00380 | Chloroplast encoded ribosomal protein S4 (AtcpRPS4) | Ribosomes | 🡫 |  |
| AtCg00330 | 30S chloroplast ribosomal protein S14 (AtcpRPS14) | Ribosomes | 🡫 |  |
| AtCg00820 | 6.8-kDa protein of the small ribosomal subunit (AtcpRPS19) | Ribosomes | 🡫 |  |
| AtCg00900 | Encodes a chloroplast ribosomal protein S7 (AtcpRPS7) | Ribosomes | 🡫 |  |
| At5g54600 | Component uL24C of plastidial ribosome (AtcpRPL24) | Ribosomes | 🡫 |  |
| At3g52150 | Component cS22 of small subunit of plastidial ribosome (AtPSRP2) | Ribosomes | 🡫 |  |
| At5g23890 | GPI-anchored adhesin-like protein | Unknown | 🡫 |  |
| At5g27560 | DUF1995 domain protein | Unknown | 🡫 |  |
| At1g12250 | Pentapeptide protein | Unknown | 🡫 |  |
| At2g48070 | Encodes a chloroplast protein RPH1 | Unknown | 🡫 |  |
| At3g61870 | Membrane protein (ECM1) located in chloroplast | Unknown | 🡫 |  |
| At3g63540 | Thylakoid lumen PsbP-type protein of unknown function (AtPPD9) | Unknown | 🡫 |  |
| At4g02725 | Protein of unknown function | Unknown | 🡫 |  |
| At5g45170 | Protein of unknown function | Unknown | 🡫 |  |
| At2g21530 | Protein of unknown function | Unknown | 🡫 |  |
| At3g11560 | Membrane protein of unknown function | Unknown | 🡫 |  |
| At3g56140 | DUF399 family protein (AtRER6) | Unknown | 🡫 |  |
| At5g42070 | Hypothetical protein | Unknown | 🡫 |  |
| At5g24690 | Membrane protein of unknown function | Unknown | 🡫 |  |
| At5g16810 | Protein kinase superfamily protein | Unknown | 🡫 |  |
| At5g03880 | Protein of unknown function | Unknown | 🡫 |  |
| At4g20130 | PEP complex component (AtPTAC14) | Transcription/translation | 🡫 |  |
| At1g06190 | Encodes a novel ribonucleic acid-binding (AtRHON1) | Transcription/translation | 🡫 |  |
| At1g09340 | Encodes CHLOROPLAST RNA BINDING (AtCRB) | Transcription/translation | 🡫 |  |
| At2g43560 | FKBP-type protein folding catalyst (AtFKBP16-3) | Transcription/translation | 🡫 |  |
| At5g04810 | Plastidial RNA splicing factor (AtPPR4) | Transcription/translation | 🡫 |  |
| At1g69200 | Carbohydrate kinase involved in transcriptional activity (AtFLN2) | Transcription/translation | 🡫 |  |
| At2g34640 | Putative PEP complex & phytochrome signalling (AtHMR) | Transcription/translation | 🡫 |  |
| At5g13650 | Encodes SVR3, a putative chloroplast TypA elongation factor | Transcription/translation | 🡫 |  |
| At2g24060 | Putative plastidial translation initiation factor (AtIF3-2) | Transcription/translation | 🡫 |  |
| At4g20360 | Nuclear transcribed, plastid localized elongation factor (AtTufA) | Transcription/translation | 🡫 |  |
| AtCg00170 | PEP complex component (AtRpoC2) | Transcription/translation | 🡫 |  |
| At2g36390 | Encodes a starch branching enzyme (AtBE3) | Starch | 🡫 |  |
| At5g66530 | Putative aldose 1-epimerase | Starch | 🡫 |  |
| At1g56050 | GTP-binding protein-like protein (AtEngD-2) | Starch | 🡫 |  |
| At4g00490 | Encodes a chloroplast beta-amylase (AtBAM2) | Starch | 🡫 |  |
| At5g48300 | Small subunit of ADP-glucose pyrophosphorylase (AtAPS1) | Starch | 🡫 |  |
| At5g19220 | Large subunit of ADP-glucose pyrophosphorylase (ATAPL1) | Starch | 🡫 |  |
| At4g39970 | [Putative xylulose-1,5-bisphosphate phosphatase (AtCbbYb)](javascript:openRefGene('58001');markCell('0');) | Starch | 🡫 |  |
| At4g24620 | Glucose-6-phosphate isomerase (AtPGI1) | Starch | 🡫 |  |
| At2g39080 | Putative (rice PHD1)-like UDP-glucose epimerase | Starch | 🡫 |  |
| At1g69830 | Encodes a plastid-localized amylase (AtAMY3) | Starch | 🡫 |  |
| At5g11880 | Putative diaminopimelate decarboxylase (AtDAPDC2) | Starch | 🡫 |  |
| At3g26060 | [Putative atypical 2-Cys peroxiredoxin (AtPrxQ)](javascript:openRefGene('54769');markCell('0');) | Redox | 🡫 |  |
| At1g77490 | Thylakoid ascorbate peroxidase (AttAPX) | Redox | 🡫 |  |
| At5g08410 | Ferredoxin/thioredoxin reductase subunit A (AtFTRa2) | Redox | 🡫 |  |
| At4g08390 | Stromal ascorbate peroxidase (AtsAPX) | Redox | 🡫 |  |
| At3g54660 | Glutathione reductase (AtGR2) | Redox | 🡫 |  |
| At1g03680 | Stromal m-type thioredoxin (AtTrx-m1) | Redox | 🡫 |  |
| At2g25080 | Glutathione peroxidase (AtGPX1) | Redox | 🡫 |  |
| AtCg01000 | AtYcf1.1/AtTic214 | Translocation | 🡫 |  |
| At2g24020 | Suppressors of tic40 (AtSTIC2) | Translocation | 🡫 |  |
| At4g01800 | AtSECa1 | Translocation | 🡫 |  |
| At3g10350 | AtGET3B | Translocation | 🡫 |  |
| At4g23940 | AtYcf2 (AtFtsHi1/AtARC1) | Translocation | 🡫 |  |
| At5g02940 | AtPollux-L1 | Translocation | 🡫 |  |
| At5g12860 | [2-oxoglutarate/malate translocator (AtDiT1/AtpOMT1)](javascript:openRefGene('7962');markCell('0');) | Translocation | 🡫 |  |
| At4g33760 | [Putative aspartate-tRNA ligase, OKINA KUKI (AtSYDM/AtOKI1)](javascript:openRefGene('57694');markCell('0');) | tRNA | 🡫 |  |
| At5g49030 | [Putative isoleucine-tRNA ligase (AtSYIM/AtOVA2)](javascript:openRefGene('68619');markCell('0');) | tRNA | 🡫 |  |
| At1g58290 | [Putative glutamyl-tRNA reductase (AtHEMA1)](javascript:openRefGene('15399');markCell('0');) | tRNA | 🡫 |  |
| At3g48110 | [Putative organellar glycyl-tRNA synthetase (AtEDD1)](javascript:openRefGene('64381');markCell('0');) | tRNA | 🡫 |  |
| At2g04842 | Putative threonine-tRNA ligase (AtThrRS2) | tRNA | 🡫 |  |
| At5g16715 | Putative valine-tRNA ligase, (AtSYVM2) | tRNA | 🡫 |  |
| At1g67090 | Rubisco small subunit 1A (AtRBCS-1A) | Dark reaction | 🡫 |  |
| At5g38430 | Rubisco small subunit 1B (AtRBCS-1B) | Dark reaction | 🡫 |  |
| At5g38410 | Rubisco small subunit 3B (AtRBCS-3B) | Dark reaction | 🡫 |  |
| AtCg00490 | Large subunit of RUBISCO (AtRBCL) | Dark reaction | 🡫 |  |
| At3g60750 | Transketolase involved in carbon fixation (AtTKL1) | Dark reaction | 🡫 |  |
| At1g80380 | [Putative D-glycerate 3-kinase involved in photorespiration (AtGLYK)](javascript:openRefGene('25123');markCell('0');) | Dark reaction | 🡫 |  |
| At1g09130 | Putative [Non-proteolytic Clp-type protease core complex (AtClpR3)](javascript:openRefGene('63954');markCell('0');) | Proteases | 🡫 |  |
| At5g05740 | [Putative (SITE-2/S2P)-like metalloprotease (AtEGY2)](javascript:openRefGene('535');markCell('0');) | Proteases | 🡫 |  |
| At2g03390 | Putative adaptor component of Clp protease machinery (AtClpF) | Proteases | 🡫 |  |
| At1g63770 | Putative broad substrate specificity aminopeptidase (AtMPA1) | Proteases | 🡫 |  |
| At3g05350 | Putative aminopeptidase | Proteases | 🡫 |  |
| At4g25100 | [Iron superoxide dismutase (AtFSD1)](javascript:openRefGene('67693');markCell('0');) | Fe-S | 🡫 |  |
| At2g42220 | Sulfurtransferase (AtSTR9) | Fe-S | 🡫 |  |
| At2g38270 | [Glutaredoxin of iron-sulfur cluster maturation (AtCXIP2)](javascript:openRefGene('17963');markCell('0');) | Fe-S | 🡫 |  |
| At5g16660 | [Fatty acid synthesis regulator (AtCTI3)](javascript:openRefGene('52377');markCell('0');) | Fatty acid | 🡫 |  |
| At5g46290 | Beta-ketoacyl ACP synthase I (KAS1) | Fatty acid | 🡫 |  |
| At1g65260 | AtVIPP1 | Membranes | 🡫 |  |
| At1g03160 | [Chloroplast membrane fusion chaperone (AtFZL)](javascript:openRefGene('13207');markCell('0');) | Membranes | 🡫 |  |
| At3g61470 | AtLHCA2 | Light reaction |  | 🡩 |
| At3g47470 | AtLHCA4 | Light reaction |  | 🡩 |
| At1g61520 | AtLHCA3.1 | Light reaction |  | 🡩 |
| At1g29930 | AtLHCB1.3 | Light reaction |  | 🡩 |
| At2g34420 | AtLHCB1.5 | Light reaction |  | 🡩 |
| At2g05100 | AtLHCB2.1 | Light reaction |  | 🡩 |
| At5g54270 | AtLHCB3.1 | Light reaction |  | 🡩 |
| At3g08940 | AtLHCB4.2 | Light reaction |  | 🡩 |
| At2g40100 | AtLHCB4.3 | Light reaction |  | 🡩 |
| At4g10340 | AtLHCB5 (AtCP26) | Light reaction |  | 🡩 |
| At1g15820 | AtLHCB6 (AtCP24) | Light reaction |  | 🡩 |
| AtCg00340 | AtPsaB | Light reaction |  | 🡩 |
| At4g02770 | AtPsaD1 | Light reaction |  | 🡩 |
| At4g12800 | AtPsaL | Light reaction |  | 🡩 |
| At2g46820 | AtPsaP | Light reaction |  | 🡩 |
| AtCg00680 | AtPsbB (AtCP47) | Light reaction |  | 🡩 |
| AtCg00280 | AtPsbC (AtCP43) | Light reaction |  | 🡩 |
| AtCg00270 | AtPsbD (D2) | Light reaction |  | 🡩 |
| AtCg00710 | AtPsbH | Light reaction |  | 🡩 |
| AtCg00580 | PSII cytochrome b559 (AtPsbE) | Light reaction |  | 🡩 |
| AtCg00540 | Putative apocytochrome cytochrome b_6_f component (AtPetA) | Light reaction |  | 🡩 |
| At4g03280 | [Putative of cytochrome b_6_f component (AtPetC)](javascript:openRefGene('64425');markCell('0');) | Light reaction |  | 🡩 |
| At1g44575 | AtPsbS (AtNPQ4) | Light reaction |  | 🡩 |
| At4g09650 | Encodes chloroplast ATPase delta-subunit (AtcpATP-delta) | Light reaction |  | 🡩 |
| AtCg00120 | Encodes chloroplast ATPase alpha-subunit (AtcpATP-alpha) | Light reaction |  | 🡩 |
| AtCg00480 | Encodes chloroplast ATPase beta-subunit (AtcpATP-beta) | Light reaction |  | 🡩 |
| At1g67350 | Plant-specific component of NADH dehydrogenase complex (AtP1) | Light reaction |  | 🡩 |
| At2g27730 | Plant-specific component of NADH dehydrogenase complex (AtP2) | Light reaction |  | 🡩 |
| AtCg01110 | Putative component of subcomplex A of chloroplast NDH (AtNdhH) | Light reaction |  | 🡩 |
| AtCg00420 | Putative component of subcomplex A of chloroplast NDH (AtNdhJ) | Light reaction |  | 🡩 |
| At4g37925 | [Putative component of subcomplex A of chloroplast NDH (AtNdhM)](javascript:openRefGene('69714');markCell('0');) | Light reaction |  | 🡩 |
| At2g39470 | [Putative of lumenal subcomplex of chloroplast NDH (AtPnsL1](javascript:openRefGene('64232');markCell('0');)) | Light reaction |  | 🡩 |
| At1g14150 | [Putative of lumenal subcomplex of chloroplast NDH (AtPnsL2](javascript:openRefGene('63332');markCell('0');)) | Light reaction |  | 🡩 |
| At4g39710 | [Putative component of chloroplast NDH (AtPnsL4](javascript:openRefGene('69021');markCell('0');)) | Light reaction |  | 🡩 |
| At4g31390 | [Kinase required for plastoquinone homeostasis (AtPGR6)](javascript:openRefGene('57573');markCell('0');) | Light reaction |  | 🡩 |
| At1g74470 | Putative geranylgeranyl reductase (AtGGR) | Light reaction |  | 🡩 |
| At1g54570 | [Putative phytyl ester synthase (AtPES1)](javascript:openRefGene('65415');markCell('0');) | Light reaction |  | 🡩 |
| At4g01050 | [Putative ferredoxin NADP-oxidoreductase protein (AtSTR4)](javascript:openRefGene('36326');markCell('0');) | Light reaction |  | 🡩 |
| At4g19170 | [Putative carotenoid cleavage dioxygenase (AtNCED4/AtCCD4)](javascript:openRefGene('56998');markCell('0');) | Light reaction |  | 🡩 |
| At1g08550 | [Putative violaxanthin de-epoxidase (AtVDE1/AtNPQ1)](javascript:openRefGene('60011');markCell('0');) | Light reaction |  | 🡩 |
| At5g58140 | Phototropin2 (AtPHOT2) | Light reaction |  | 🡩 |
| At1g71810 | Putative ABC1-type atypical kinase (AtABC1K5) | Other |  | 🡩 |
| At1g21440 | Putative carboxy-phosphonoenolpyruvate phosphonomutase | Other |  | 🡩 |
| At3g09580 | Putative flavin containing amine oxidoreductase | Other |  | 🡩 |
| At2g34460 | Putative NAD-dependent epimerase/dehydratase | Other |  | 🡩 |
| At5g05200 | Putative ABC1-type atypical kinase (AtABC1K9) | Other |  | 🡩 |
| At1g06690 | Putative NAD(P)-dependent aldo-keto reductase | Other |  | 🡩 |
| At4g25900 | Putative aldose 1-epimerase | Other |  | 🡩 |
| At1g50450 | Putative saccharopine dehydrogenase | Other |  | 🡩 |
| At1g33590 | Putative PNP signalling peptide receptor guanylyl cyclase | Other |  | 🡩 |
| At4g39980 | [Putative -3deoxy-D-arabino-heptulosonate 7-phosphate (AtDHS1)](javascript:openRefGene('63745');markCell('0');) | Other |  | 🡩 |
| At5g63400 | [putative adenylate kinase (AtAMK4](javascript:openRefGene('44387');markCell('0');)) | Other |  | 🡩 |
| At3g25760 | [Putative allene oxide cyclase (AtERD12)](javascript:openRefGene('54746');markCell('0');) | Other |  | 🡩 |
| At5g42650 | [Putative allene oxide synthase (AtAOS)](javascript:openRefGene('59766');markCell('0');) | Other |  | 🡩 |
| At3g14210 | [Putative GDSL-type carboxylesterase (AtGLP63/)](javascript:openRefGene('60766');markCell('0');) | Other |  | 🡩 |
| At4g26530 | Putative fructose-bisphosphate aldolase (AtFBA5) | Other |  | 🡩 |
| At4g09000 | [Putative 14-3-3 signalling regulator protein (AtGRF1)](javascript:openRefGene('56509');markCell('0');) | Other |  | 🡩 |
| At4g01900 | [Putative regulatory nitrogen sensor protein (AtGLB1)](javascript:openRefGene('56222');markCell('0');) | Other |  | 🡩 |
| At1g68010 | [Putative hydroxypyruvate reductase (AtHPR1)](javascript:openRefGene('63424');markCell('0');) | Other |  | 🡩 |
| At3g44310 | Putative indole-3-acetonitrile nitrilase (AtNIT1) | Other |  | 🡩 |
| At4g11010 | Putative nucleoside diphosphate kinase (AtNDPK3) | Other |  | 🡩 |
| At4g19410 | [Putative pectin acetylesterase (AtPAE7)](javascript:openRefGene('23874');markCell('0');) | Other |  | 🡩 |
| At1g20620 | Putative transnitrosylase involved in nitric oxide signalling, (AtCAT3) | Other |  | 🡩 |
| At1g10760 | Putative alpha-glucan water dikinase (AtSEX1) | Other |  | 🡩 |
| At4g02510 | AtTOC159 | Translocation |  | 🡩 |
| At3g46740 | AtTOC 75-III | Translocation |  | 🡩 |
| At3g17970 | AtTOC64-III | Translocation |  | 🡩 |
| At4g23430 | AtTIC32 | Translocation |  | 🡩 |
| At2g24820 | AtTIC55 | Translocation |  | 🡩 |
| At5g22640 | AtTic100 | Translocation |  | 🡩 |
| At1g76405 | Outer envelope pore 21B-like protein (AtOEP21-L) | Translocation |  | 🡩 |
| At2g17695 | Outer envelope protein AtOEP23 | Translocation |  | 🡩 |
| At5g42960 | Outer envelope pore AtOEP24B-like protein | Translocation |  | 🡩 |
| At3g48870 | [Clp-type protease complex (AtHSP93-III)](javascript:openRefGene('67087');markCell('0');) | Translocation |  | 🡩 |
| At1g65410 | ABC-type component of lipid transfer complex (AtTGD3) | Translocation |  | 🡩 |
| At5g64290 | [Putative glutamate:malate translocator (AtpDCT1)](javascript:openRefGene('229');markCell('0');) | Translocation |  | 🡩 |
| At2g46910 | Putative FIBRILLIN plastoglobule-associated protein (AtPAP10) | Starch |  | 🡩 |
| At4g04020 | Putative [FIBRILLIN plastoglobule-associated protein (AtPAP1)](javascript:openRefGene('67370');markCell('0');) | Starch |  | 🡩 |
| At4g22240 | [Putative FIBRILLIN plastoglobule-associated protein (AtPAP2)](javascript:openRefGene('67657');markCell('0');) | Starch |  | 🡩 |
| At3g23400 | [Putative FIBRILLIN plastoglobule-associated protein (AtPAP6)](javascript:openRefGene('54633');markCell('0');) | Starch |  | 🡩 |
| At1g51110 | Putative FIBRILLIN plastoglobule-associated protein (AtPAP12) | Starch |  | 🡩 |
| At3g26070 | Putative FIBRILLIN plastoglobule-associated protein (AtPAP4) | Starch |  | 🡩 |
| At5g28840 | Putative GDP-mannose epimerase (AtGME) | Starch |  | 🡩 |
| At1g32900 | Putative granule-bound starch synthase (AtGBSS) | Starch |  | 🡩 |
| At2g01140 | [Putative fructose-bisphosphate aldolase (AtFBA3)](javascript:openRefGene('16445');markCell('0');) | Starch |  | 🡩 |
| At2g41950 | Protein of unknown function | Unknown |  | 🡩 |
| At3g20820 | Protein of unknown function | Unknown |  | 🡩 |
| At3g28220 | Protein of unknown function | Unknown |  | 🡩 |
| At1g16790 | Protein of unknown function | Unknown |  | 🡩 |
| At5g37360 | Protein of unknown function | Unknown |  | 🡩 |
| At1g76180 | [LEA class-2 protein of unknown function (AtERD14)](javascript:openRefGene('16200');markCell('0');) | Unknown |  | 🡩 |
| At3g56650 | Unknown function thylakoid lumen PsbP-type protein (AtPPD6) | Unknown |  | 🡩 |
| At3g27925 | Encodes a DegP protease (AtDegP) | Proteases |  | 🡩 |
| At1g66670 | [Putative component of Clp-type protease complex (AtClpP3)](javascript:openRefGene('64084');markCell('0');) | Proteases |  | 🡩 |
| At3g54400 | Putative aspartyl protease protein | Proteases |  | 🡩 |
| At1g73990 | [Putative SppA-like protease (AtSppA)](javascript:openRefGene('65699');markCell('0');) | Proteases |  | 🡩 |
| At3g13920 | Putative translation initiation factor (AtTIF4A-1) | Transcription/translation |  | 🡩 |
| At1g07930 | [Putative eEF1-alpha translation elongation factor (AteEF1A2)](javascript:openRefGene('13439');markCell('0');) | Transcription/translation |  | 🡩 |
| At3g16000 | [Putative plastid transcriptional activity regulation protein (AtMFP1)](javascript:openRefGene('68980');markCell('0');) | Transcription/translation |  | 🡩 |
| At1g80030 | [Plastidial Hsp40/DnaJ molecular co-chaperone (AtDJA7)](javascript:openRefGene('65787');markCell('0');) | Chaperone/assembly |  | 🡩 |
| At5g14910 | [Auxiliary plastidial ribosome biogenesis factor (AtCRASS)](javascript:openRefGene('58669');markCell('0');) | Chaperone/assembly |  | 🡩 |
| At4g09010 | [Putative plastidial peroxidase-like protein (AtAPX4)](javascript:openRefGene('56510');markCell('0');) | Redox |  | 🡩 |
| At5g17170 | [Putative rubredoxin-like protein (AtENH1)](javascript:openRefGene('40848');markCell('0');) | Redox |  | 🡩 |
| At3g08920 | Putative sulfurtransferase (AtSTR10) | Fe-S |  | 🡩 |
| At3g59780 | Putative inactive sulfurtransferase-like protein (AtSTR19) | Fe-S |  | 🡩 |
| At3g15690 | Putative regulator of acetyl-CoA carboxylase complex (AtBADC3) | Fatty acids |  | 🡩 |
| At3g45140 | Putative 13-lipoxygenase (AtLOX2) | Fatty acids |  | 🡩 |
| At1g72150 | [Putative PATELLIN membrane trafficking protein (AtPATL1)](javascript:openRefGene('16020');markCell('0');) | Membranes |  | 🡩 |

| **Table S2. Genotyping primers.**  This table supports figure 4 of the manuscript and significantly contributes to the transparency and reproducibility of genotyping experiments. | | |
| --- | --- | --- |
| **Gene** | **Allele** | **Primer sequences** |
| *GET3B* | Wild type | GCACCGCTTCTTTGTATAGC |
|  |  | AGAGAGTAGCCTTAGCGTATGA |
|  | SALK_017702C (*get3b-2*) | GCGTGGACCGCTTGCTGCAACT |
|  |  | AGAGAGTAGCCTTAGCGTATGA |
|  | ProUBI10:GET3B transgenes | ATTGCCAATTTTCAGCTCCA |
|  |  | TTCGAACCTTGAAGCGAGTT |
| *SRP54* | Wild type | ATATTGTTGGCTGGGCTCCA |
|  |  | TGAAGCCTCCCTGCAGTATC |
|  | WiscDsLox289_292B14 (*srp54-3*) | ATATTGTTGGCTGGGCTCCA |
|  |  | AACGTCCGCAATGTGTTATTAAGTTGTC |
| *ALB4* | Wild type | TGCTGGAATAAATCCCCTTG |
|  |  | CATCCGAGAGGTGGGTGT |
|  | SALK136199(*alb4-1*) | TGCTGGAATAAATCCCCTTG |
|  |  | ATTTTGCCGATTTCGGAAC |
| *STIC2* | Wild type | GCTTCCGGTTTTATCTTCTCG |
|  |  | GAACACGTACAGCTTCCACTTG |
|  | SALK_001500 (*stic2-3*) | GCTTCCGGTTTTATCTTCTCG |
|  |  | GCGTGGACCGCTTGCTGCAACT |
|  | WiscDsLox445D01 (*stic2-4*) | GCTTCCGGTTTTATCTTCTCG |
|  |  | AACGTCCGCAATGTGTTATTAAGTTGTC |
